# Gymnosperm and fern multifunctional bHLH transcription factors control stomatal development

**DOI:** 10.1101/2025.11.10.687565

**Authors:** Miki Zaizen-Iida, Kaotar Elhazzime, Aleksia Fanni Maria Vaattovaara, Vojtech Schmidt, Sofie Johansson, Thomas Dobrenel, Steffen Grebe, Ulrika Egertsdotter, Ove Nilsson, Anne Vatén

**Affiliations:** Department of Organismal and Evolutionary Biology and Department of Molecular and Integrative Biosciences, Faculty of Biological and Environmental Sciences and Viikki Plant Science Centre, University of Helsinki, Helsinki, Finland; Umeå Plant Science Centre (UPSC), Department of Forest Genetics and Plant Physiology, Swedish University of Agricultural Sciences, 901 83 Umeå, Sweden; Institute for Atmospheric and Earth System Research (INAR)/Forest Sciences, Faculty of Agriculture and Forestry and Viikki Plant Science Center, University of Helsinki, Helsinki, Finland; Renewable Bioproducts Institute, Georgia Institute of Technology, Atlanta, GA, USA

**Keywords:** stomatal development, fern, gymnosperm, cross-species analysis, stomatal evolution, evolution of developmental regulation, stomatal pore formation, guard cell maturation

## Abstract

Stomatal development is driven by a conserved family of basic Helix-loop-helix (bHLH) transcription factors. In the angiosperm model *Arabidopsis thaliana*, *SPEECHLESS* (*SPCH*), *MUTE* and *FAMA* show pronounced yet incompletely separated stage-specific functions. How the functional specialization observed within angiosperms evolved has remained unclear. Here, we perform the first functional analysis of stomatal regulators from the fern *Ceratopteris richardii* and the gymnosperm *Picea abies* (Norway spruce). We show that both species possess a single *SPCH/MUTE*-like gene (*CrSM* and *PaSM*) together with a single *FAMA*-like gene (*CrFAMA* and *PaFAMA*). Cross-species complementation in *Arabidopsis* revealed broad but biased developmental capacities: all four bHLHs could promote stomatal-lineage entry, whereas SM more effectively supported early-to-intermediate transitions and FAMA showed stronger capacities at later stages. *PaSM* and *PaFAMA* showed overlapping but distinct expression patterns within spruce stomatal lineage, with *PaSM* enriched at early to middle developmental stages and *PaFAMA* present across a broader developmental window. Dominant-negative perturbation of either gene in spruce was associated with altered stomatal-file patterning while producing partially distinct effects on mature stomatal-complex morphology. Together, these findings show that PaSM and PaFAMA display broad and in-part overlapping developmental roles within spruce stomatal lineage. Our data supports the model in which angiosperm stage specialization emerged through progressive partitioning of broader, overlapping developmental capacities.

## Introduction

Stomata are microscopic epidermal pores that mediate CO₂ uptake for photosynthesis while limiting transpirational water loss, and their acquisition was likely a key innovation enabling the colonization of land by plants^1^. In most land plants, the basic stomatal structure is conserved and consists of a pair of guard cells (GCs) surrounding a stomatal pore^2,3^. In some lineages, however, this core structure is accompanied by additional specialized cells termed subsidiary cells (SCs)^4^. For example, grasses develop SCs that facilitate GC movements and contribute to the distinctive morphology and function of grass stomatal complexes^5^. Despite such morphological variation, the core regulatory machinery underlying stomatal development appears to be widely conserved.

Stomatal development is controlled by bHLH transcription factors of the subgroup Ia (bHLH1a). In angiosperm model plant, *Arabidopsis thaliana*, three genes encoding bHLH transcription factors *SPCH*, *MUTE*, and *FAMA* act sequentially, together with the broadly expressed bHLH partners ICE1/SCRM and SCRM2, to control successive stomatal-lineage transitions^6–9^. *SPCH* initiates the stomatal lineage by driving asymmetric cell divisions (ACDs), *MUTE* promotes guard mother cell (GMC) fate, and *FAMA* establishes GC identity. Orthologs of these genes are broadly conserved across angiosperms and generally retain similar core functions^10–12^, although in some lineages, they have acquired additional lineage-specific roles^5,11–16^. Although *SPCH*, *MUTE* and *FAMA* have distinct stage-defining functions, their activities are not completely temporally separated. Recent work showed that *SPCH* persists into GMCs and young GCs^17^, where the relative levels and activities of *SPCH* and *FAMA* regulate GC division and expansion^18^.

How this pronounced, yet partially overlapping, stage-biased specialization evolved has remained unresolved. Phylogenetic analysis have yielded alternative reconstructions of bHLH1a gene-family evolution. In one scenario, the angiosperm stomatal bHLH module arose through duplication and subsequent diversification of an ancestral bHLH1a gene^19^. However, other reconstructions cannot exclude the possibility that early land plants already possessed multiple bHLH1a genes and that the smaller gene content within some extant lineages resulted from secondary gene loss^20^. These alternative gene-family histories do not, by themselves, reveal the ancestral functional states of the encoded proteins or the extent to which developmental functions are partitioned among coexisting paralogs outside angiosperms. Domain-swapping and mutational analysis of *Arabidopsis* stomatal bHLHs have shown how sequence divergence could generate the specialized activities of *SPCH*, *MUTE*, and *FAMA*^21^. Although these findings provide a mechanistic model for progressive functional divergence following gene duplication, they do not establish whether this was the historical route by which stomatal bHLH specialization evolved.

Functional studies in bryophytes have nevertheless provided insight into the developmental capacities of early-diverging land-plant bHLH1a. In the moss *Physcomitrium patens*, *PpSMF1* partially complements *Arabidopsis mute* and *fama*, whereas native loss of *PpSMF1* abolishes stomatal formation in the moss sporophyte^3,22^. In the astomatous liverwort *Marchantia polymorpha*, its sole bHLH1a gene functions in non-stomatal developmental contexts but retains the capacity to partially support stomatal development in *Arabidopsis*^23,24^. Together, these findings demonstrate that extant single-copy bryophyte bHLH1a proteins can possess broad developmental capacities. However, neither they distinguish between alternative histories of bHLH1a gene duplication and loss nor determine whether, under a scenario of ancestral gene multiplicity, coexisting paralogs were already functionally specialized. Thus, irrespective of gene-family history, how the pronounced yet incomplete stage-biased specialization of angiosperm stomatal bHLHs evolved has not been resolved.

Available genomes from several fern and gymnosperm species encode, or are predicted to encode, one *SPCH/MUTE*-like gene together with one *FAMA-like* gene^19,20^. This two-gene complement provides an opportunity to test whether coexisting paralogs retain broadly overlapping developmental capacities or exhibit substantial functional partitioning. Comparative functional characterization of these genes can therefore provide evidence about the degree of functional specialization among stomatal bHLHs outside angiosperms.

Here, we compare the regulatory capacities of *SPCH/MUTE-* and *FAMA-*like transcription factors from the gymnosperm *Picea abies* and the fern *Ceratopteris richardii* through cross-species complementation in *Arabidopsis*. In parallel, we tested *PaSM* and *PaFAMA* expression across spruce stomatal development and examined the effects of fusing *PaSM* and *PaFAMA* to a transcriptional repression domain in transgenic spruce. Together, these approaches enable functional comparison of evolutionary divergent bHLHs in *Arabidopsis* context while independently providing native-context evidence for PaSM- and PaFAMA-associated functions in gymnosperm stomatal development.

To our knowledge, this study provides the first comparative functional analysis of stomatal bHLH transcription factors from a fern and a gymnosperm. All four fern and gymnosperm transcription factors promoted asymmetric stomatal-lineage entry divisions in *Arabidopsis*, and PaSM additionally promoted stomatal pore formation, revealing developmental capacities that span transitions primarily controlled by distinct regulators in angiosperms. Native-context analysis further provided initial evidence for partially distinct PaSM- and PaFAMA-associated activities in spruce. Together, these findings uncover previously unrecognized broad and overlapping developmental capacities among fern and gymnosperm stomatal bHLHs, with evidence for partial functional differentiation between the two paralogs in spruce.

## Results

### Norway spruce stomatal development shows conserved and distinct features

We first investigated stomatal development in Norway spruce needles. Stomatal development proceeded from the base toward the tip of the needle and was restricted to specific cell files (Figure 1A, Supplementary Figure 1A and 1B). Spruce needles were rhombic in cross-section, with up to three stomatal cell files present on each of the four faces (Supplementary Figure 1B). We sometimes observed stomatal files shifting between adjacent epidermal cell files, suggesting some flexibility in the spatial patterning of stomatal development. To analyze early stages of stomatal development, we sampled young needles located near the vegetative meristem of spruce seedlings. Because stomatal development was largely complete in needles longer than 10 mm, we focused on needles shorter than 10 mm (Supplementary Figure 1A). In spruce stomatal files, protodermal cells underwent divisions that appeared morphologically symmetric (Figure 1A and 1B). The resulting daughter cells subsequently diverged in morphology: one elongated into a rectangular cell, whereas the other became rounded, consistent with distinct developmental trajectories (Figure 1A and 1C). This divergence was already apparent at the earliest post-division stages that we could resolve, suggesting that daughter-cell fate asymmetry is established before, during, or shortly after division. By reconstructing stomata-forming cell files from mature stomata toward younger developmental stages, we inferred that the elongated rectangular cell was most likely the GMC, as it later underwent a symmetric division, whereas the rounded daughter cell became a non-stomatal cell. Consistent with previous observations in pines^25^, these results suggest that spruce protodermal cells undergo an equal division whose daughter cells differentiate into a GMC and a polar SC, respectively. We therefore refer to the non-stomatal daughter cell in spruce as the polar SC (Figure 1A and 1C). During GMC differentiation, cells in the adjacent non-stomatal row differentiated into lateral subsidiary mother cells (SMCs) (Figure 1A and 1C). Subsequently, the GMC and lateral SMCs underwent symmetric and asymmetric divisions, respectively, producing a pair of GCs and two SCs flanking the GCs (Figure 1A and 1D). After these divisions, the GCs began to sink into the hypodermal layer (Figure 1A, 1E and 1F). In the mature stomatal complex, the GCs were fully sunken within the hypodermal layer, as reported in other conifers^25^. Together, these observations indicate that spruce stomatal development includes conserved features such as the establishment of cell fate asymmetry, differentiation and symmetric division of the GMC, and GC formation. At the same time, it also shows distinctive characteristics, including the recruitment of multiple SCs and the displacement of GCs from the epidermis into a subepidermal position.

**Figure 1.**
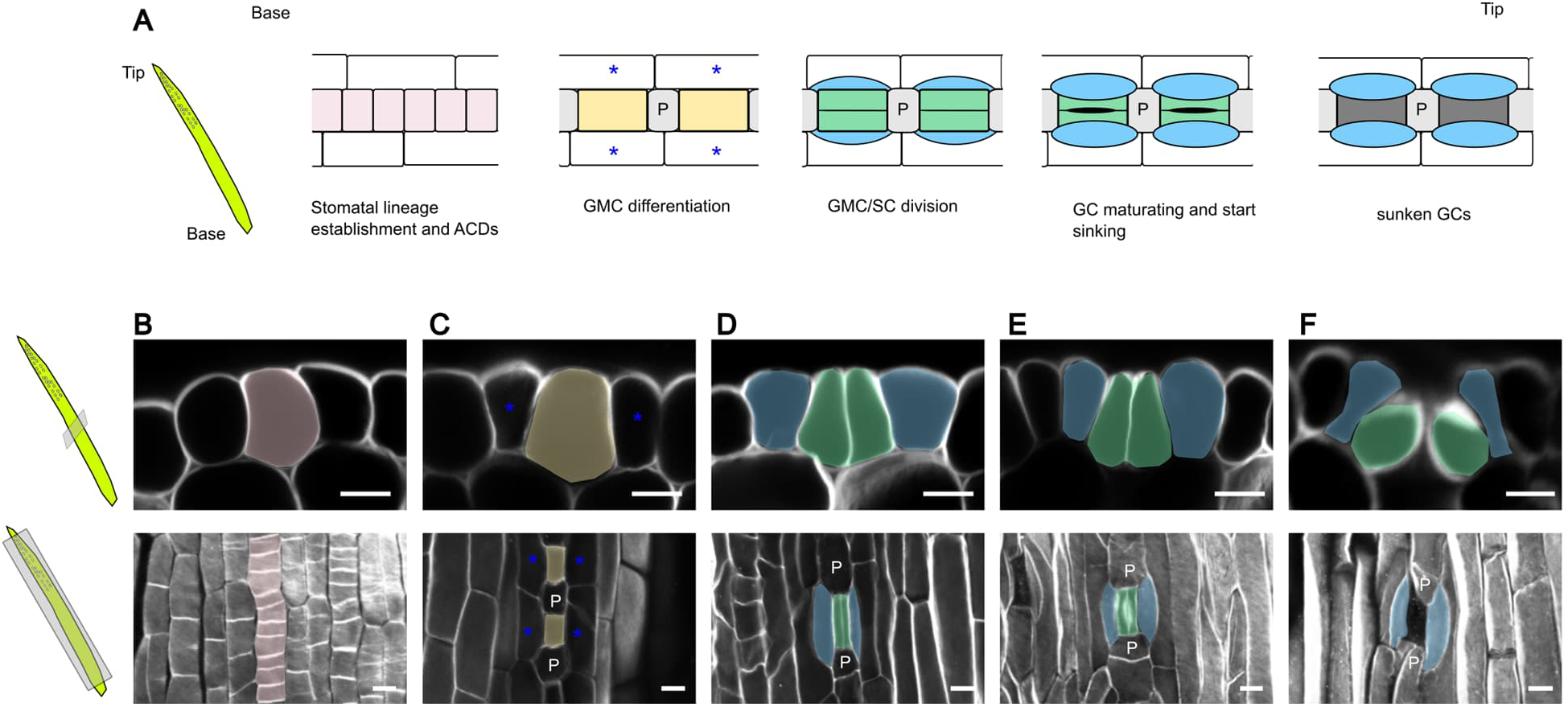
Conserved and distinct features of stomatal development in Norway spruce. **A,** Schematic representation of stomatal development in a young Norway spruce needle near the SAM. Stomatal development proceeds from the base of the needle toward the tip. **B-F,** Detailed structures of young spruce needles. **Upper panels** show cross-sections; **Lower panels** show lateral views. **B,** Symmetric division of stomatal precursor cells. **C,** The divided protodermal cells differentiate into an elongated GMC and a round polar SC. The cell adjacent to GMC subsequently develops into lateral SMC. **D,** The GMC divides symmetrically to produce a pair of GCs, while the lateral SMC divides asymmetrically to generate lateral SCs. **E,** After GMC division, GCs begin to sink into the hypodermal layer. **F,** GCs are located at the hypodermal layer. Pink, yellow, green and blue indicate stomatal precursor cell, GMC, GC, and lateral SC, respectively; in **B**, these colors are shown as semi-transparent overlays. Blue asterisk and “P” indicate lateral SMC and polar S, respectively. White, SR2200. Scale bars, 10 µm.

### Norway spruce genome contains two genes encoding bHLH1a members in SM and FAMA subclades

To identify bHLH1a coding genes in spruce, we performed a phylogenetic analysis and constructed a corresponding phylogenetic tree containing a set of species including bryophytes, a lycophyte, ferns, gymnosperms and angiosperms (Figure 2A). The phylogenetic tree resolved five subclades. The single *Marchantia polymorpha* bHLH1a gene is placed outside these subclades, near the root defined by the two *Arabidopsis* bHLH3c outgroup genes. One of the subclades consists only fern sequences. *SPCH/MUTE* (*SM*) and *FAMA* subclades form sister groups. The *FAMA* subclade included genes from all the analyzed species except *Marchantia polymorpha,* whereas *SM* subclade contains only ferns, gymnosperms and angiosperms. Gymnosperms have also subclade with conserved number of homologs closely related to angiosperm subclade including *TEC1/c* and *MYCc7/70* homologs. We identified that *Picea abies* genome assembly contains a single *SPCH/MUTE*-like gene (*PaSM*; *MA_120c02g0010*) and a *FAMA*-like gene (*PaFAMA*; *MA_57244g0010*) (Figure 2A). We next compared the domain structures of *Arabidopsis thaliana* and *Picea abies* SPCH, MUTE, and FAMA orthologs (Figure 2B). Both PaSM and PaFAMA contain the conserved bHLH1a and C-terminal domains characterizing the subgroup 1a transcription factors^26,27^. Additionally, PaSM has a truncated MITOGEN-ACTIVATED PROTEIN KINASE target domain (MAPKTD), a region important for the regulation of AtSPCH protein stability^21,28^. Because AtSPCH variants lacking MAPKTD phosphorylation sites can promote GMC differentiation, a capacity normally associated with MUTE, the truncated MAPKTD in PaSM raised the possibility that PaSM might possess both SPCH- and MUTE-like activities. However, we noticed that PaSM lacks a MUTE unique domain, which function has not been defined. PaFAMA contains putative FAMA-specific domains as well as the LxCxE motif, which is essential for the interaction with RETINOBLASTOMA-RELATED (RBR) protein in *Arabidopsis*^29^. Based on phylogenetic analysis and domain structures, we hypothesized that PaSM may functionally resemble both SPCH and MUTE or alternatively either SPCH or MUTE, whereas PaFAMA is likely to retain FAMA-like functions.

**Figure 2.**
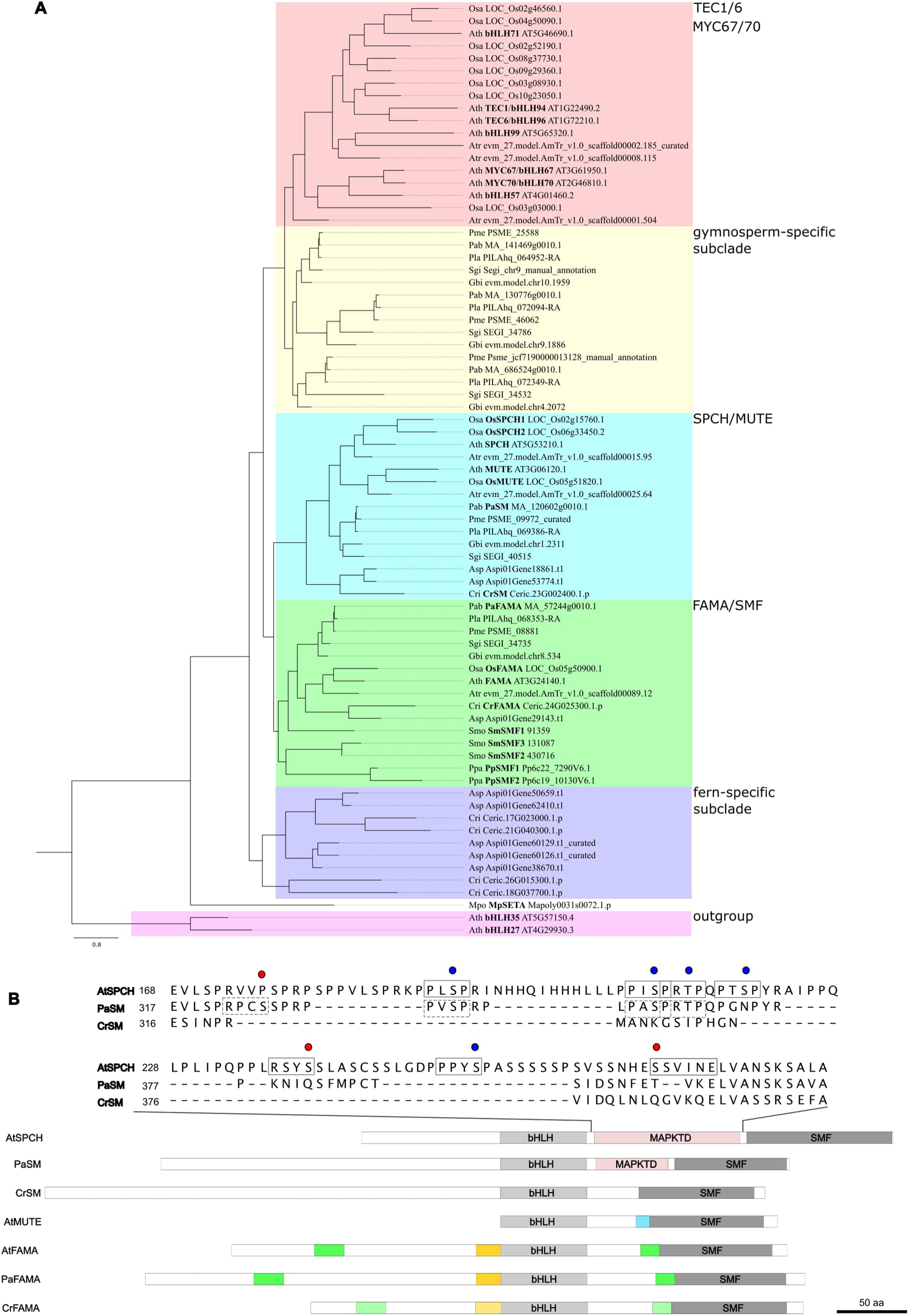
Genomes of ferns and gymnosperms contain a single *SPCH/MUTE*-like and a *FAMA*-like gene. **A,** Maximum-likelihood bHLH phylogenetic tree of subfamily Ia. Species abbreviations are as follows: Asp, *Alsophila spinulosa* (fern); Ath, *Arabidopsis thaliana* (dicot); Atr, *Amborella trichopoda* (basal angiosperm); Cri, *Ceratopteris richardii* (fern); Gbi, *Ginkgo biloba* (gymnosperm); Mpo, *Marchantia polymorpha* (liverwort); Osa, *Oryza sativa* (monocot); Pab, *Picea abies* (gymnosperm); Pla, *Pinus lambertiana* (gymnosperm); Pme, *Pseudotsuga menziesii* (gymnosperm); Ppa, *Physcomitrium patens* (moss); Sgi, *Sequoiadendron giganteum* (gymnosperm); Smo, *Selaginella moellendorffii* (lycophyte). Background color shows each subclade: purple, fern-specific subclade; green, FAMA/SMF subclade; turquoise, SPCH/MUTE (SM) subclade; yellow, gymnosperm-specific subclade; red, TEC1/6 and MYC67/70 containing subclade. *Arabidopsis* bHLH3c genes were used as outgroups and are shown in pink. **B,** Comparison of domain structures of tracheophyte bHLH1a proteins from the SM and FAMA subclades. Each bHLH1a contains bHLH domain (light gray) and a conserved C-terminal SMF domain (dark gray). AtSPCH and PaSM have MAPKTD or putative MAPKTD, shown in red, although the putative MAPKTD in PaSM is truncated. AtMUTE has MUTE specific domain (blue), whereas CrSM has neither MAPKTD nor MUTE specific domain. FAMA orthologs contain a putative FAMA specific domain (green) and a putative bHLH domain extension (yellow). For putative FAMA specific domain and putative bHLH domain extension, color intensity indicates amino acid identity relative to *Arabidopsis* sequence. From the N terminus, amino acid identities of the corresponding regions were 72.7%, 77.8% and 50.0% in PaFAMA, and 13.6%, 50.0% and 35.7% in CrFAMA. Gray rectangles are MAPK3/6 target motifs or SnRK2 motifs previously characterized in earlier studies^28,79^. Gray dashed rectangles are putative MAPK3/6 or SnRK2 target motifs predicted from amino acid sequences, based on the consensus motifs PxS/TP for MAPK3/6 and RxxS or SxxxxE for SnRK2. Blue circles indicate experimentally characterized or putative MPK3/6 phosphorylation residues. Red circles indicate experimentally characterized or putative SnRK2 phosphorylation residues.

### *PaSM* and *PaFAMA* transcripts peak in needle regions enriched for different stages of stomatal development

Our phylogenetic analysis identified PaSM and PaFAMA as candidate regulators of stomatal development in spruce. To examine whether their expression patterns vary among needle regions enriched with different phases of stomatal development, we collected basal and distal regions from needles of different size classes to generate four developmental stages enriched for successive stages of stomatal development. We then analyzed *PaSM* and *PaFAMA* transcript abundance in these regions by quantitative real-time PCR (qRT–PCR). Based on confocal microscopy-based imaging, developmental stage 1, 2, 3 were enriched predominantly for ACDs, GMC differentiation/division, and maturing stomata, respectively (Figure 3A–3C). Stage 4, collected from the distal region of longer needles, contained more mature stomata and was used to represent the latest developmental stage. We nevertheless observed some morphological variation among needles of similar size, suggesting that needle length alone does not fully reflect stomatal developmental progression.

**Figure 3.**
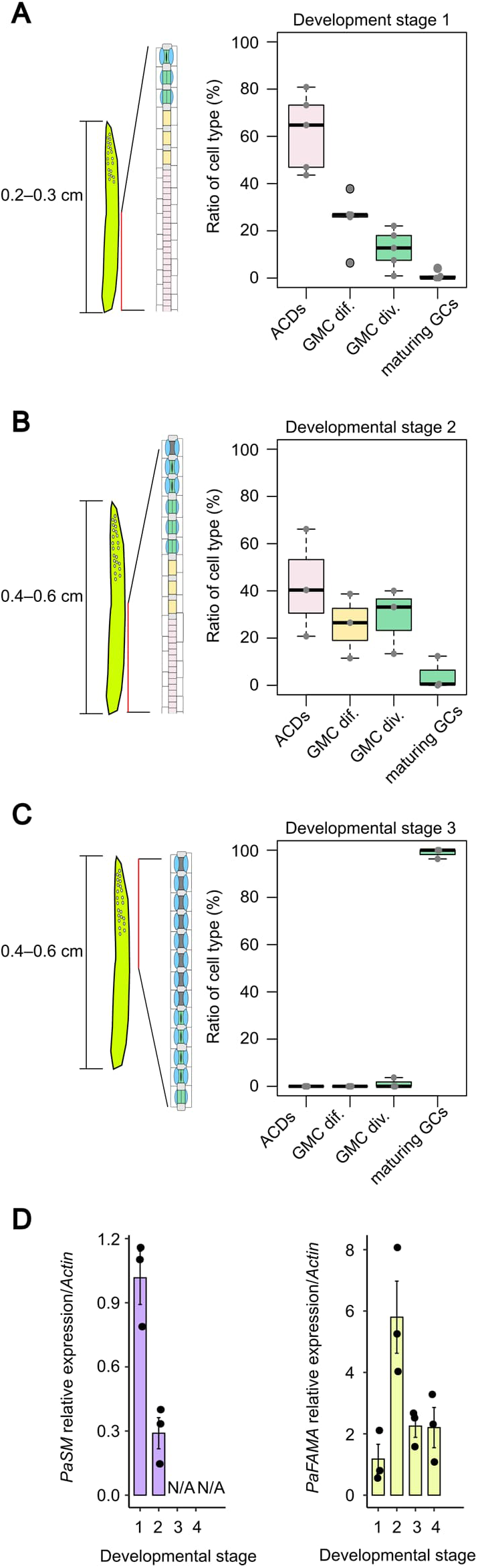
*PaSM* and *PaFAMA* show temporally distinct gene expression during Norway spruce stomatal development. **A–C,** Characterization of stomatal developmental stages in developing Norway spruce needles. Stage 1 is characterized predominantly by ACDs (**A**), stage 2 by GMC differentiation and division (**B**), and stage 3 by maturing GCs (**C**). Stage 4 represents a later stage containing more mature GCs. Schematic diagrams adjacent to the plots summarize the major stomatal developmental events represented by each stage. **D,** qRT-PCR analysis of *PaSM* and *PaFAMA* expression at each developmental stage. Expression levels were normalized to *Actin* and are shown relative to stage 1. *PaSM* was preferentially expressed at stage 1, with little or no expression at stages 3 and 4, which are characterized by maturing and mature stomata. *PaFAMA* was predominantly expressed at stage 2, with comparable expression levels at stages 1, 3, and 4. Bars indicate mean relative expression, and error bars indicate ± s.e.m. Individual points represent biological replicates.

We next quantified *PaSM* and *PaFAMA* transcript levels across stages 1–4. *PaSM* expression was highest in the ACD-enriched stage and decreased markedly in the GMC differentiation/division-enriched stage. *PaSM* transcripts were not detected in the stages enriched for maturing and mature stomata, as no Ct values were obtained (Figure 3D). These results indicate that *PaSM* is associated primarily with early stomatal lineage stages and was most abundant in regions enriched for ACDs.

By contrast, *PaFAMA* expression showed a broader developmental distribution. Its transcript levels were highest in the GMC differentiation/division-enriched stage. However, *PaFAMA* transcripts were also clearly detectable in the ACD-enriched stage and remained detectable in the later stages enriched with maturing and more mature GCs stages (Figure 3D). Although expression in the later stages showed some variation among samples, these data indicate that *PaFAMA* expression persists well beyond the GMC-enriched stage. This broader pattern suggests that *PaFAMA* is associated most strongly with GMC differentiation and division, while transcripts remained detectable in regions enriched for both earlier and later stages of stomatal development.

For defining spatial expression patterns for *PaSM* and *PaFAMA*, we performed HCR RNA-FISH analysis of young Norway spruce needles and found that *PaSM* and *PaFAMA* transcripts are indeed present within the stomatal lineage. *PaSM* puncta were observed in cells resulting from ACDs and GMCs ongoing differentiation, whereas no clear signal was observed in GMCs at the later division stage in the sections examined (Supplementary Figure 2A). *PaFAMA* puncta were observed across a broader range of developmental stages, including ACDs, GMC differentiation, and GMC division, and were occasionally detected also in mature GCs (Supplementary Figure 2B). These observations align with the qRT–PCR results.

Together, these results reveal that *PaSM* and *PaFAMA* shows distinct yet partially overlapping expression patterns in stomatal cell lineage. Considered alongside their sequence features (Figure 2B), these expression patterns prompted us to test whether *PaSM* possesses *SPCH*- and/or *MUTE*-like regulatory capacities and whether *PaFAMA* retains *FAMA*-like activity.

### *PaSM* and *PaFAMA* can promote early stomatal-lineage transitions in *Arabidopsis*

To assess the developmental capacities of *PaSM* and *PaFAMA* in *Arabidopsis*, we introduced *PaSM* and *PaFAMA* with triple yellow fluorescence protein (Venus) fusion in the *Arabidopsis spch-3*, *mute-1*, and *fama-1* mutants under the control of the respective *AtSPCH*, *AtMUTE*, or *AtFAMA* promoters and assessed their ability to restore stomata production and to rescue seedling lethality. This approach allowed us to test whether these proteins could support stomatal-lineage transitions controlled by *AtSPCH*, *AtMUTE* or *AtFAMA* in *Arabidopsis*.

Expression of *pAtSPCH:PaSM* in *spch*-3, which normally produces an epidermis composed only of pavement cells and display seedling lethality^6^, resulted in frequent cell divisions, production of irregularly shaped stomata and occasional single GCs (Figure 4A). Moreover, *pAtSPCH:PaSM* was sufficient to rescue seedling lethality and recover seed production. However, we noticed that these lines displayed cracks between epidermal cells and occasionally partial detachment of the epidermis from inner layers (Supplementary Figure 3A), which prevented accurate quantification of stomatal number in mature cotyledons collected from 10-day-old seedlings. To mitigate this issue, we analyzed younger cotyledons from 5-day-old seedlings, which exhibited fewer cracks, and confirmed that *pAtSPCH:PaSM* expression restored some extent of stomata production in the absence of AtSPCH (Figure 4B). These results indicate that *PaSM* supported entry into the stomatal lineage, although it does not fully restore the wild-type epidermal phenotype.

**Figure 4.**
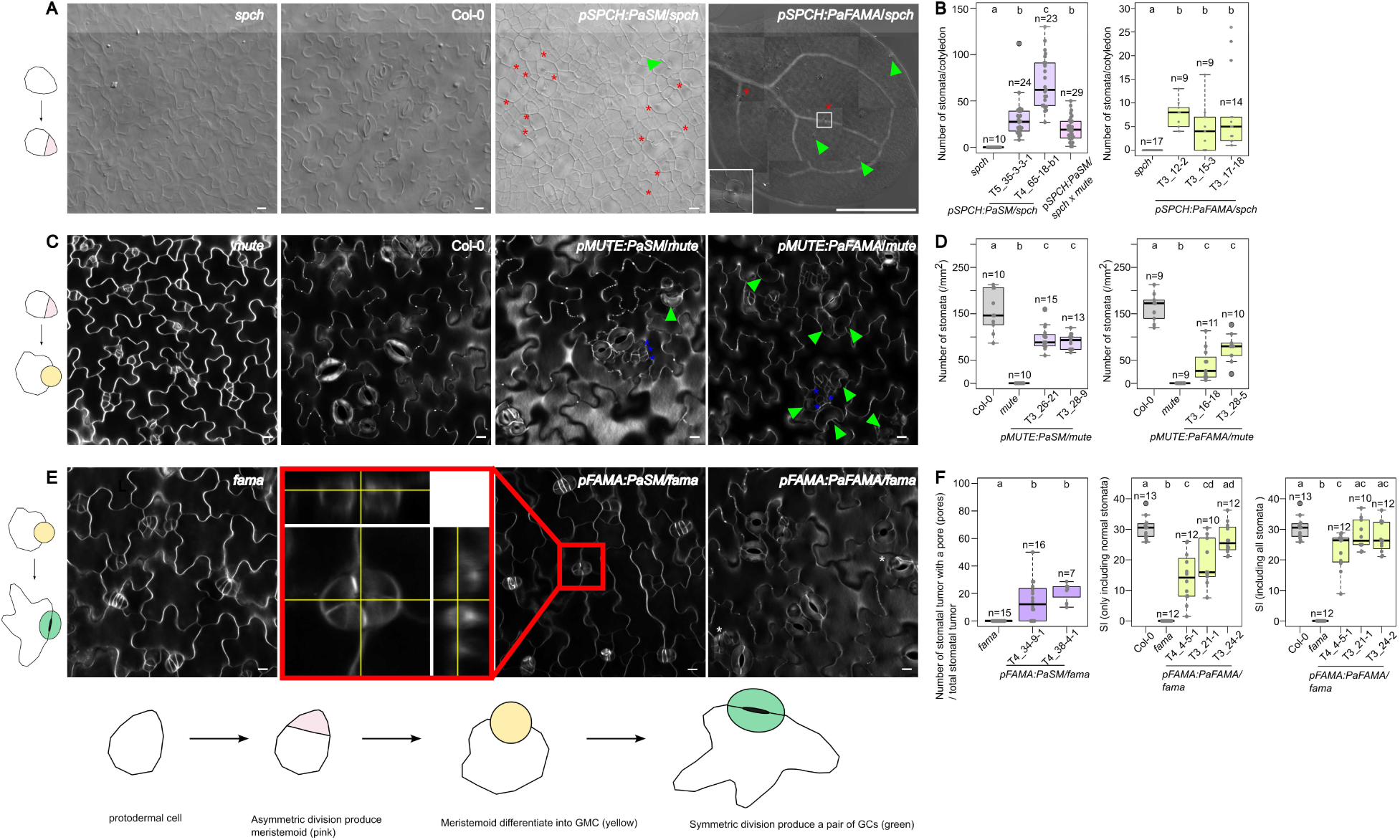
Both *PaSM* and *PaFAMA* complement multiple steps of *Arabidopsis* stomatal development. Cross-species complementation analysis using *Arabidopsis thaliana*. The schematic illustrates the progression of stomatal development from early lineage entry to mature guard cell formation. **A,** DIC images of 10-day-old abaxial cotyledons of Col-0, *spch-3* mutant, *pAtSPCH:PaSM/spch-3*, and *pAtSPCH:PaFAMA/spch-3*. **B,** Number of stomata per abaxial cotyledon in 5-day-old *pAtSPCH:PaSM/spch-3* seedlings and 10-day-old *pAtSPCH:PaFAMA/spch-3* seedlings. **C,** Confocal images of 10-day-old abaxial cotyledons of Col-0, *mute-1* mutant, *pAtMUTE:PaSM/mute-1*, and *pAtMUTE:PaFAMA/mute-1*. **D,** SD in 10-day-old abaxial cotyledons of *pAtMUTE:PaSM/mute-1* and *pAtMUTE:PaFAMA/mute-1*. **E,** Confocal images of 10-day-old abaxial cotyledons of *fama-1* mutant, *pAtFAMA:PaSM/fama-1*, and *pAtFAMA:PaFAMA/fama-1*. The red box indicates the region enlarged in the inset in *pAtFAMA:PaSM/fama-1*. An optical section from the z-stack reveals a pore-like structure within the stomatal tumor. **F,** Number of stomatal tumors having pores per total stomatal tumor in abaxial cotyledon in 10-day-old *pAtFAMA:PaSM/fama-1* seedlings. SI on the abaxial side of cotyledons in 10-day-old *pAtFAMA:PaFAMA/fama-1* seedlings. SI was calculated using either normal stomata only or both normal stomata and “abnormal stomata”. For **B, D,** and **F,** statistical significance was assessed by the Steel-Dwass-Critchlow-Fligner test; different letters indicate statistically significant differences between groups (*p* < 0.05). Box plots show the median and interquartile range, with whiskers extending to values within 1.5× the interquartile range. Individual data points are shown as beeswarm plots. Cell walls were stained with SR2200 (**C**,**E**).Red asterisks, green triangles, blue asterisks, and white asterisks indicate stomata with aberrant morphology, single GCs, small cells and “abnormal”, respectively. Scale bars, 500 µm in *pAtSPCH:PaFAMA/spch-3* in **A** and 10 µm in all other images in **A, C,** and **E**.

Next, we tested whether PaFAMA could support stomatal-lineage entry when expressed under the *AtSPCH* promoter in the *spch* mutant. p*AtSPCH*:*PaFAMA* expression was sufficient to drive formation of stomata and stomatal lineage cells in the cotyledons of the *spch-3*, albeit infrequently (Figure 4A and 4B). The seedling-lethal phenotype of *spch-3* was occasionally rescued; however, these plant lines failed to produce seeds. We observed substantial variability in stomatal numbers both across the cotyledons and between individual seedlings. To obtain more accurate view of this phenotype, we therefore counted the total number of stomata per cotyledon. Similar to *pAtSPCH:PaSM/spch, pAtSPCH:PaFAMA/spch* seedlings also showed an epidermal peeling-off phenotype (Supplementary Figure 3A). In most cases, however, this phenotype was less severe than that observed in *pAtSPCH:PaSM/spch,* enabling us to quantify stomatal numbers in 10-day-old seedlings. Our analysis showed that three independent lines with *pAtSPCH:PaFAMA* consistently induced stomata production in the *spch-3* (Figure 4B), indicating that PaFAMA can support stomatal lineage entry.

Because the unique function of AtSPCH among group 1a bHLHs is to promote ACDs, we next examined whether PaSM and PaFAMA can drive ACDs in the absence of AtSPCH. We observed abaxial epidermis of young cotyledons in the *pAtSPCH:PaSM/spch-3* and *pAtSPCH:PaFAMA/spch-3* lines and found that some of the divisions were indeed morphologically asymmetric (Supplementary Figure 3B). These findings indicate that both PaSM and PaFAMA can support stomatal-lineage entry and induce occasional morphologically asymmetric divisions in the *spch* background. The presence of stomata in these lines suggests that at least some morphologically asymmetric divisions may have been accompanied by fate asymmetry. These two genes show different ability to rescue seedling lethality, suggesting a functional and/or structural difference in the stomatal complexes produced within these lines.

We then introduced *PaSM* and *PaFAMA* into *mute-1* mutant. While both *pAtMUTE:PaSM/mute-1* and *pAtMUTE:PaFAMA/mute-1* lines showed wild-type like stomata, they still retained some arrested meristemoids similar to *mute* mutant^7^. Additionally, we occasionally observed small cells with unknown identity (Figure 4C). In *pAtMUTE:PaFAMA/mute-1*, we frequently found single GCs, which were rarely observed in *pAtMUTE:PaSM/mute-1* (Supplementary Fig. 3C). We compared stomatal density (SD) between the wild type, *mute-1*, and *mute-1* complemented by *pAtMUTE:PaSM* or *pAtMUTE:PaFAMA*. Two independent lines of both *pAtMUTE:PaSM/mute-1* and *pAtMUTE:PaFAMA/mute-1* showed significantly increased SD compared to the *mute* mutant, although SD remained lower than that of the wild type (Figure 4D). We defined relative proportions of stomatal cell types and found that *pAtMUTE:PaSM/mute-1* showed higher relative number of stomata, whereas *pAtMUTE:PaFAMA/mute-1* displayed higher relative numbers of meristemoids and single GCs (Supplementary Figure 3C). Both transgenes partially rescued the defects in seedling viability and seed production observed in *mute*. These results indicate that both PaSM and PaFAMA can partially support AtMUTE function in *Arabidopsis*. However, PaSM promoted production of a higher proportion of stomata relative to meristemoids than PaFAMA did, consistent with a greater capacity to promote progression through MUTE-dependent developmental steps, including stages of GMC differentiation and symmetric division.

To further test whether PaSM act as a multifunctional transcription factor spanning both SPCH- and MUTE-associated steps, we introduced *PaSM* into *spch-3 mute-1* double mutant under the control of *AtSPCH* promoter. Notably, the *AtMUTE* promoter is active in a subset of the broader stomatal-lineage domain covered by the *AtSPCH* promoter^6,30^. The resulting plants showed similar phenotype as *pAtSPCH:PaSM/spch-3* lines, which showed stomata with aberrant morphology and cracks between epidermal cells (Supplementary Figure 3A). We noticed that *pAtSPCH:PaSM/spch-3 mute-1* displayed fewer number of stomata per cotyledon than *pAtSPCH:PaSM/spch-3* (Figure 4B). However, stomata formation in the absence of both *AtSPCH* and *AtMUTE* indicates that PaSM has ability to partially support both stomatal-lineage entry and subsequent MUTE-dependent progression to stomata formation when expressed in *Arabidopsis*. The broad developmental capacity of PaSM raises the possibility that overlapping SPCH- and MUTE-like activities can coexist within a single SPCH/MUTE-like transcription factor.

### *PaSM* promotes pore-like structures whereas *PaFAMA* partially restores GC differentiation in *Arabidopsis*

We explored whether PaSM and PaFAMA can drive the last cell fate transition in the *Arabidopsis* stomatal lineage and support the function of AtFAMA. Expression of *PaSM* in *fama-1* under the control of the *AtFAMA* promoter produced viable seedlings with a partially rescued aspects of the *fama* phenotype, although the plants later failed to produce seeds. We found clusters of small tumor-like structures resembling caterpillar-like clusters of cells present in the *fama*^8^; however, their division orientation was altered compared to typical *fama* tumors. We also occasionally found stomata-like structures; they were composed of GC-like cells with additional divisions, consistent with incomplete maintenance of terminal GC differentiation (Figure 4E). We classified both *fama* tumor-like and stomata-like cells as “stomatal tumors” and quantified the proportion exhibiting pore-like characteristics. In *fama* mutant, tumors did not exhibit pore-like characteristics, whereas 15–20% of stomatal tumors in the *pAtFAMA:PaSM*/*fama-1* lines formed pore-like structures (Figure 4F). These findings indicate that PaSM can promote the appearance of pore-like structures within a subset of *fama* tumors but does not restore stable terminal GC differentiation in *Arabidopsis*.

In the *pAtFAMA:PaFAMA/fama-1* lines, we observed both normal stomata and “abnormal stomata”, which was characterized by additional cell divisions within GCs (Figure 4E). We first quantified the Stomatal Index (SI) including only normal stomata; one of the three independent lines exhibited SI similar to that of the wild type. When both normal stomata and “abnormal stomata” were included, two out of the three lines displayed SI values that were not significantly different from wild type (Figure 4F). “Abnormal stomata” showed diverse structures; most abundant defects resembled the stomata-in-stomata phenotype^29^. Additional variants included: (i) parallel divisions along the stomatal pore axis within GCs, (ii) both parallel and vertical divisions in each GC within the same pair of GCs, and (iii) vertical divisions followed by extra divisions of the smaller daughter cell, producing structures reminiscent of *fama*-like tumors (Supplementary Figure 3D and 3E). These data indicate that PaFAMA can partially substitute for AtFAMA in *Arabidopsis* by supporting stomatal pore formation and guard cell differentiation. However, the frequent additional divisions indicate incomplete enforcement of terminal cell-cycle arrest and are consistent with PaFAMA retaining residual capacity to promote cell divisions at late stomatal-lineage stages.

PaSM and PaFAMA possess broad capacity in stomatal development, albeit with a degree of functional bias; PaSM primarily rescued early-to-middle developmental stages, whereas PaFAMA predominantly rescued middle-to-late stages. This implies that these bHLH transcription factors may operate with intrinsic functional preferences.

### C.fern *SM* and *FAMA* display overlapping stomatal regulatory capacities in *Arabidopsis*

Our phylogenetic analysis showed that both SM and FAMA orthologs are also present in ferns (Figure 2A and 2B). To place the gymnosperm findings in a broader evolutionary context, we investigated two bHLH1a transcription factors from the fern *Ceratopteris richardii*, CrSM and CrFAMA, and asked whether they exhibit overlapping or specialized developmental capacities. We tested their activities through cross-species complementation in *Arabidopsis* stomatal mutants.

Expression of *CrSM* or *CrFAMA* under the *AtSPCH* promoter in the *spch-3* background restored seedling viability and seed production, and induced stomatal-lineage cell formation and occasional ACDs. This indicates that both *CrSM* and *CrFAMA* can support entry into the stomatal-lineage in *Arabidopsis*. However, their effects differed: CrSM frequently produced excessive stomata or stomatal clusters, whereas CrFAMA primarily generated abnormally differentiated cells and only infrequently produced stomata. (Figure 5A, 5B, Supplementary Figure 4A and 4B).

**Figure 5.**
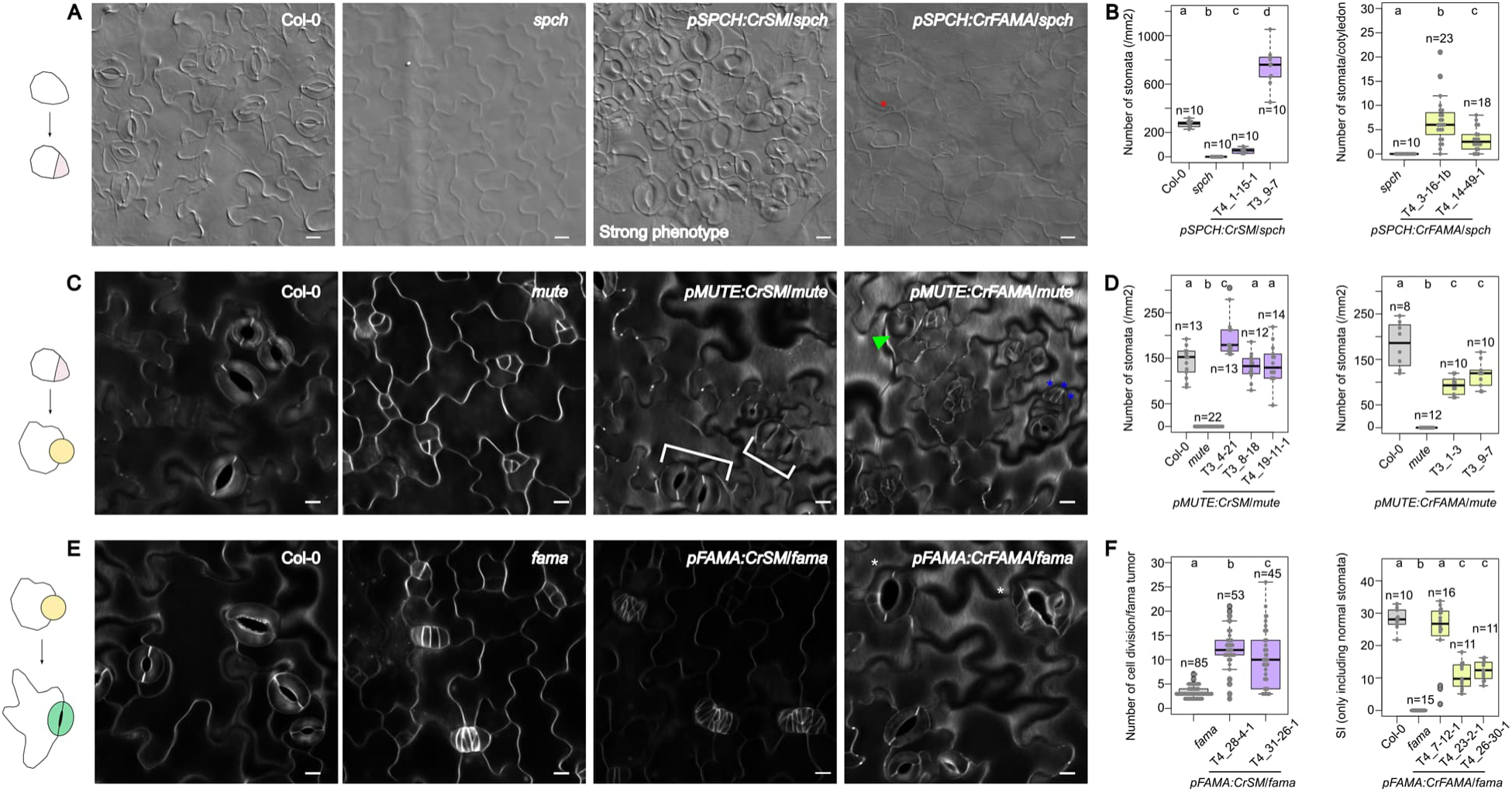
Both *CrSM* and *CrFAMA* show complementation capacity similar to Norway spruce stomatal genes. **A**, DIC images of 10-day-old abaxial cotyledons of Col-0, *spch-3* mutant, *pAtSPCH:CrSM/spch-3*, and *pAtSPCH:CrFAMA/spch-3*. *pAtSPCH:CrSM/spch-3* showed a phenotype resembling *AtMUTE* overexpression^7^. **B**, SD in 5-day-old abaxial cotyledons of *pAtSPCH:CrSM/spch-3.* Number of stomata in 5-day-old abaxial cotyledons of *pAtSPCH:CrFAMA/spch-3.* **C,** Confocal images of 10-day-old abaxial cotyledons of Col-0, *mute-1* mutant, *pAtMUTE:CrSM/mute-1*, and *pAtMUTE:CrFAMA/mute-1.* **D**, Stomatal density in 10-day-old abaxial cotyledons of either *pAtMUTE:CrSM/mute-1* or *pAtMUTE:CrFAMA/mute-1.* **E,** Confocal images of 10-day-old abaxial cotyledons of Col-0, *fama-1* mutant, *pAtFAMA:CrSM/fama-1*, and *pAtFAMA:CrFAMA/fama-1.* **F,** Number of cell division per *fama* tumor in 10-day-old adaxial cotyledons of *pAtFAMA:CrSM/fama-1*. SI of abaxial cotyledon in 10-day-old *pAtFAMA:CrFAMA/fama-1* seedlings. Only normal stomata were quantified. For **B, D,** and **F,** statistical significance was assessed by the Steel-Dwass-Critchlow-Fligner test; different letters indicate statistically significant differences between groups (*p* < 0.05). Box plots show the median and interquartile range, with whiskers extending to values within 1.5× the interquartile range. Individual data points are shown as beeswarm plots. Cell walls were stained with SR2200 (**C**,**E**). Red asterisks, white bracket, green triangles, blue asterisks, and white asterisks indicate stomata with aberrant morphology, pairs of stomata, single GCs, small cells and “abnormal stomata”, respectively. Scale bars,10 µm in **A, C,** and **E**.

Both *CrSM* and *CrFAMA* also partially complemented *mute-1* when expressed under the *AtMUTE* promoter and restored seedling viability and seed production, indicating that they can support MUTE-associated developmental transitions in *Arabidopsis*. *CrSM* lines produced paired stomata and Venus-tagged CrSM was detected in young GCs, where the endogenous AtMUTE is typically not expressed. The persistence of CrSM protein into this later developmental stage may lead to an additional round of symmetric division in young GCs (Supplementary Figure 4D and 4E). In *CrSM* lines, SD was comparable to or higher than wild type (Figure 5B), whereas *CrFAMA* resulted in a weaker rescue with mixed stomatal-lineage phenotypes (Figure 5C, 5D and Supplementary Figure 4F).

When expressed under the *AtFAMA* promoter, *CrFAMA*, but not *CrSM*, partially complemented *fama-1*. CrFAMA produced both normal and “abnormal stomata”, with phenotypes similar to those observed in *pAtFAMA-PaFAMA/fama-1* lines, whereas CrSM failed to restore stomatal differentiation and instead enhanced further cell proliferation within *fama*-like tumors (Figure 5E, 5F and Supplementary Figure 4G–J).

Together, these results suggest that C.fern SM and FAMA show overlapping activities during early and middle stomatal development, whereas only the CrFAMA display the capacity to support FAMA-associated terminal GC differentiation when expressed in *Arabidopsis*. C.fern data also supports the idea that bHLH transcription factors operate with intrinsic functional preferences.

### Perturbation of PaSM and PaFAMA produces overlapping and distinct effects on spruce stomatal development

Although our cross-species complementation analysis revealed the developmental capacities of stomatal bHLHs in *Arabidopsis*, their native roles and the extent of functional overlap remain unclear. To investigate the function of PaSM and PaFAMA in the native context, we generated transgenic spruce lines constitutively expressing chimeric repressor domain SRDX^31^ fused with PaSM and PaFAMA under the control of 35S promoter, hereafter referred to as *PaSM-SRDX* and *PaFAMA-SRDX*. Transgene expression was confirmed by qRT-PCR using proembryogenic mass samples (S5A). Transgenic lines from both constructs germinated and produced viable seedlings resembling the control lines (Figure 6A and Supplementary Figure 5B).

**Figure 6.**
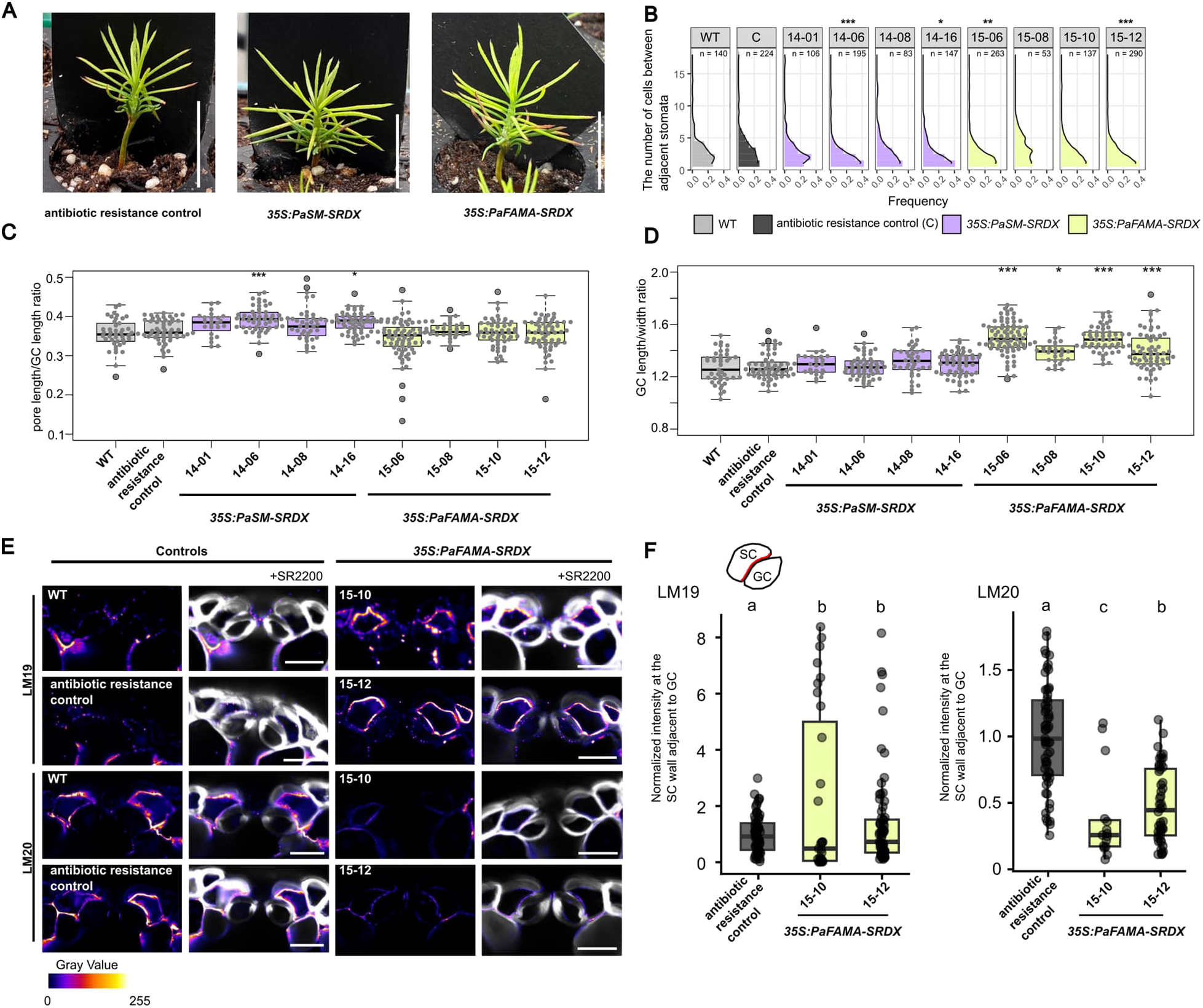
*PaSM* and *PaFAMA* display overlapping and distinct function in stomatal development. **A**, Approximately one-month-old seedling images of antibiotic-resistant control, *35S:PaSM-SRDX* and *35S:PaFAMA-SRDX* lines. **B**, Density plots show the frequency distribution of the number of cells between adjacent stomata. **C**, Pore length/GC length ratio of wild type, antibiotic-resistant control, *35S:PaSM-SRDX* and *35S:PaFAMA-SRDX* lines. **D**, GC length/width ratio of wild type, antibiotic-resistant control, *35S:PaSM-SRDX* and *35S:PaFAMA-SRDX* lines. **E**, Immunolabeling images of wild type, antibiotic-resistant control, *35S:PaSM-SRDX* and *35S:PaFAMA-SRDX* lines. LM19 and LM20 were used to detect distinct HG epitopes. **F**, Quantification of LM19 and LM20 signal intensity in the SC wall at the interface with GC. n≥20 subsidiary cells per line, from three independent experiments using three different plants. For **B–D,** Statistical significance was assessed using the Kruskal–Wallis test followed by Dunn’s multiple comparisons test against the antibiotic-resistant control. Asterisks indicate significant differences compared with the antibiotic - resistant control: *P < 0.05, **P < 0.01, ***P < 0.001. Box plots show the median and interquartile range, with whiskers extending to values within 1.5× the interquartile range. Individual data points are shown as beeswarm plots. For **f**, Statistical significance was assessed using a linear mixed-effects model (F-test for the fixed effect of line; LM19: P=0.005, LM20: P=9.76 x10^-7^) followed by pairwise comparisons (Tukey method, no additional adjustments). Different letters indicate statistically significant differences between groups (*P* < 0.05). Scale bars, 1 cm in **A**, 20 µm in **E**.

We analyzed mature needles collected from approximately three-month-old transgenic and control plants. All the lines formed stomata, suggesting either functional redundancy with other putative regulators of stomata production or reduced efficiency of SRDX system in spruce. To assess whether perturbation of PaSM and PaFAMA affected the cell division behavior in stomatal files, we quantified the number of cells between adjacent stomata. In both *PaSM-SRDX* and *PaFAMA-SRDX* lines, two out of four independent transgenic lines showed a significant reduction in the number of cells locating between stomata compared with the antibiotic resistance control line (Figure 6B). The reduced number of intervening cells may reflect changes in cell proliferation or patterning within stomatal files. However, analysis of mature needles cannot distinguish whether there are changes within stomatal-lineage initiation events, division of intervening cells.

To determine whether perturbation of PaSM and PaFAMA was associated with changes in mature stomatal-complexes, we performed detailed characterization of stomatal complex morphology in control, *PaSM-SRDX* and *PaFAMA-SRDX* lines. Strikingly, two out of four transgenic lines with *PaSM-SRDX* showed distinct differences in GC and stomatal complex morphology with a larger pore length/GC length ratio compared with antibiotic resistance control line (Figure 6C). This indicates that *PaSM-SRDX* expression can alter stomatal pore geometry relative to GC length. In addition, the GC length/width ratio was consistently increased in the all four *PaFAMA-SRDX* lines compared to the antibiotic resistance control line (Figure 6D), suggesting that *PaFAMA* contributes to the regulation of GC shape, possibly by promoting lateral expansion.

Because GC shape is strongly influenced by cell wall properties, we next examined whether this morphological phenotype was associated with altered pattern of pectin epitopes in the stomatal complex. We analyzed labeling with LM19 and LM20 antibodies, which preferentially recognize homogalacturonan (HG) epitopes associated with lower and higher degrees of methyl esterification, respectively. In *PaFAMA-SRDX* lines, LM20 labeling was significantly reduced, whereas LM19 labeling was significantly increased at the SC wall adjacent to GCs in two out of four independent lines. This reciprocal labeling patterns suggest that there is a relative shift in detectable HG epitopes towards lower methyl-esterification within this cell wall domain.

Thus, PaFAMA perturbation is associated with altered HG epitope pattern at the GC–SC interface. However, because GC shape was altered in all the four *PaFAMA-SRDX* lines whereas significant changes in LM19 and LM20 labeling were detected in only two lines, altered HG epitope patterning alone cannot fully explain the GC shape phenotype. Notably, *PaSM-SRDX* lines did not show altered GC shape or HG epitope patterning, distinguishing the effects of the two constructs on mature stomatal complexes (Figure 6E, 6F and Supplementary Figure 5D).

In addition to changes in mature stomatal morphology, we examined whether the GMC division pattern was affected. In both *PaSM-SRDX* and *PaFAMA-SRDX* lines, we occasionally observed cells containing oblique or perpendicular division walls within stomatal files (Supplementary Figure 5C). These cells were more rounded than surrounding non-stomatal epidermal cells, suggesting that they may represent GMC-like cells with abnormal division orientation. The rarity of this phenotype was consistent with the narrow developmental window during which GMC divisions occur and because of this, we were not able to quantify the phenotype reliably. Nevertheless, this observation suggests that PaSM and PaFAMA activity may contribute to the proper cell division plane orientation within GMCs.

Taken together, perturbation of PaSM and PaFAMA produced partially overlapping and distinct effects on spruce stomatal development. Both *PaSM-SRDX* and *PaFAMA-SRDX* were associated with fewer epidermal cells separating adjacent stomata, indicating altered cell division patterns of stomatal cell files. The effects on mature stomatal complexes were distinct; *PaFAMA-SRDX* altered GC shape (4/4 lines) and was associated (2/4 lines) with altered HG epitope patterning at SC wall adjacent to GC, whereas *PaSM-SRDX* altered stomatal pore geometry relative to GC length (2/4 lines) without detectable changes in GC shape or HG labeling. Together with their developmental expression patterns, these results support the involvement of both PaSM and PaFAMA in spruce stomatal development while indicating partially distinct effects on stomatal-complex formation.

## Discussion

We functionally characterized SPCH/MUTE- and FAMA-like bHLH1a transcription factors from Norway spruce and C. fern and found that these regulators combine broad developmental capacities with distinct functional biases. All four bHLHs could support multiple transitions of the *Arabidopsis* stomatal lineage, including stomatal-lineage entry, a capacity that is strongly associated with SPCH in angiosperms. At the same time, their activities were not equivalent: SM more effectively supported early-to-intermediate transitions, whereas FAMA showed stronger capacities during intermediate-to-late development. In spruce, *PaSM* and *PaFAMA* also showed overlapping but distinct developmental expression patterns and produced overlapping but partially distinct phenotypes when perturbed. Together, these findings indicate partial functional partitioning: broad developmental capacities are retained despite biases in the transitions and, in spruce, partial divergence in developmental deployment and native phenotypic effects. This pattern is consistent with a model in which the pronounced stage specialization of angiosperm stomatal regulators emerged through progressive partitioning of broader developmental capacities.

Previous work showed that AtMUTE has only a limited ability to compensate for loss of AtSPCH function^32^, consistent with functional divergence between SPCH and MUTE. In contrast, PaSM partially complemented both *spch* and *mute* mutants and supported stomata formation in the *spch mute* double-mutant background. These findings suggest PaSM possesses broad developmental capacities spanning from stomatal-lineage entry to the subsequent MUTE-associated progression. The excessive number of cell divisions and aberrant stomatal structures observed in the *pAtSPCH:PaSM/spch-3* lines resembled phenotypes produced by an AtSPCH variant with disrupted MAPK-dependent negative regulation^28^. This is consistent with the presence of only a partial MAPKTD in PaSM and raises the possibility that PaSM is regulated differently from AtSPCH with respect to protein stability. At the same time, some high-stringency MAPK target sites implicated in SPCH regulation are present in PaSM^28^, suggesting that elements of this regulatory mechanism may predate the full elaboration of the angiosperm *SPCH* module. Notably, *PaSM* did not rescue the *fama* mutant, however, PaSM supported pore formation within tumor-like stomatal structures in the *fama* background, a phenotype not reported for AtSPCH/*fama* complementation lines^3^. Thus, it remains possible that developmental capacity of PaSM in *Arabidopsis* extends beyond that associated with AtSPCH.

*PaFAMA*, by contrast, partially rescued the *fama* and *mute* mutant. This pattern is broadly consistent with previous reports showing that AtFAMA and BdFAMA can rescue both *mute* and *fama* mutants^12^, and with studies demonstrating that the *Marchantia* ortholog MpSETA and the moss ortholog *PpSMF1* can partially rescue *Arabidopsis mute* and *fama* mutants^3,23^. Unexpectedly, PaFAMA also partially supported stomatal-lineage entry in the *spch* background, a capacity not reported for AtFAMA, MpSETA, or PpSMF1^3,12,23^. This suggests that besides supporting a conserved late-lineage role similar to AtFAMA, PaFAMA may also possess a broader developmental capacity than previously recognized for FAMA-like transcription factors. Together, these results show that both PaSM and PaFAMA can act across multiple stomatal-lineage transitions in *Arabidopsis*, although their functional emphases differ: PaSM can more effectively support early and intermediate transitions, whereas PaFAMA is more effective in promotion of terminal GC differentiation.

Comparison of the gymnosperm and fern transcription factors places the broad developmental capacities observed for PaSM and PaFAMA in a wider evolutionary context. Cross-species complementation assays showed that the two bHLH1a transcription factors from C.fern also function across multiple stages of the *Arabidopsis* stomatal lineage. CrSM partially complemented both *spch* and *mute* mutants, indicating that, like PaSM, it can support stomatal-lineage entry and subsequent MUTE-associated progression. Whereas PaSM contains a truncated MAPKTD, CrSM lacks this domain entirely (Figure 2B). The *pAtSPCH:CrSM/spch* lines formed stomatal clusters resembling those produced by constitutively active AtSPCH variants lacking the MAPKTD^28^. This phenotypic similarity raises the possibility that the absence of a MAPKTD contributes to altered regulation of CrSM activity in *Arabidopsis*. Both PaSM and CrSM also retain an N-terminal region that is absent from AtMUTE, which cannot substitute effectively for AtSPCH^6^. Although this correlation does not establish a causal role, it raises the possibility that the N-terminal region contributes to SPCH-like activity. Unlike PaSM, CrSM failed to rescue *fama* and instead enhanced cell proliferation within *fama* tumors. This resembles the phenotype caused by ectopic expression of *CYCD5;1* under the control of *AtMUTE* promoter in *fama-1* mutants^33^. Because AtFAMA normally contributes to CYCD5;1 repression, one possible interpretation is that CrSM cannot impose the FAMA-dependent cell-cycle arrest required for terminal GC differentiation. Alternatively, CrSM may retain an active capacity to promote the expression of cell-cycle regulators. Also CrFAMA displayed a broad activity profile: it partially complemented *spch, mute*, and *fama* mutants, thus demonstrating capacities spanning early, intermediate, and terminal stages of stomatal development. Together, these findings show that broad and overlapping developmental capacities are not restricted to the Norway spruce bHLHs. Both fern bHLHs functioned across more than one stomatal-lineage transition in *Arabidopsis*, but their functional emphases remained distinct: CrSM predominantly supported early and intermediate development.

The additional divisions in GCs observed in *pAtFAMA:PaFAMA/fama-1* and *pAtFAMA:CrFAMA/fama-1* could reflect incomplete maintenance of the terminal cell-cycle arrest normally imposed by AtFAMA. However, an alternative interpretation is that PaFAMA and CrFAMA possess division-promoting activity. Recent work showed that SPCH persists at low levels into late GMCs and young GCs and that elevated *SPCH* expression in the FAMA domain induces ectopic symmetric divisions without reverting cells to an early stomatal-lineage identity^18^. The resemblance of this phenotype to the additional divisions induced by PaFAMA and CrFAMA, together with their capacities to support earlier stomatal-lineage transitions, raises the possibility that these FAMA-like transcription factors possess an SPCH-like capacity to promote cell division when expressed during late stomatal development.

Native-context analysis in spruce was consistent with partial, rather than complete, functional partitioning. *PaSM* and *PaFAMA* showed distinct but overlapping distribution of their transctips based on qRT-PCR. *PaSM* expression was highest in regions enriched for early stomatal-lineage divisions and declined strongly before late stomatal maturation. *PaFAMA* showed a broader profile, with maximal expression in regions enriched for GMC differentiation and division but detectable transcripts also across earlier and later developmental stages. By using HCR RNA-FISH, *PaSM* was similarly detected primarily in early to middle stomatal-lineage cells, whereas *PaFAMA* occurred across a broader range of stages. These expression patterns indicate partial partitioning of developmental deployment between the two genes, even though their proteins retain overlapping capacities when tested in *Arabidopsis*.

The effects of PaSM-SRDX and PaFAMA-SRDX further support this interpretation. Perturbation of either gene was associated in subsets of independent lines with fewer epidermal cells between adjacent stomata, suggesting partially overlapping effects on stomatal-file proliferation or patterning. Their effects on mature stomatal complexes, however, differed. PaSM-SRDX altered pore length relative to GC length, whereas PaFAMA-SRDX reproducibly altered GC shape and, in a subset of lines, HG epitope patterning at the GC–SC interface. The PaSM-SRDX pore phenotype may result indirectly from perturbation of an early developmental process; in *Arabidopsis*, polarity established during early stomatal development has been linked to specification of the future pore-forming domain^34^. By contrast, the broader developmental distribution of *PaFAMA* is consistent with its effects on mature GC morphology and cell-wall properties. These native phenotypes therefore suggest that overlapping regulatory activities coexist with partially distinct contributions to stomatal-complex development.

The pattern of *PaFAMA* transcript abundance observed here differs from the predominantly late-GMC to early-GC expression reported for *AtFAMA* and *BdFAMA*^8,12,35^. Notably, *PaFAMA* transcripts remained detectable in stage 4 samples based on the qRT-PCR data (Figure 3D), in which mature stomata predominate, raising the possibility that *PaFAMA* expression extends further into stomatal maturation than in these angiosperm systems. Our HCR RNA-FISH data provide partial support for this interpretation, as *PaFAMA* transcripts were occasionally detected in mature GCs (Supplementary Figure 2B). However, *PaFAMA* transcripts were also detected in subset of mesophyll cells (Supplementary Figure 2C), indicating that the late-stage qRT-PCR signal cannot be attributed exclusively to mature stomata. The mesophyll expression further raises the possibility that PaFAMA has functions outside the stomatal lineage. Consistent with this idea, *Arabidopsis* FAMA functions not only in GC differentiation but also in the differentiation of myrosin cells^35^, demonstrating that AtFAMA activity is not restricted to the stomatal lineage. Whether PaFAMA similarly contributes to the differentiation or function of non-stomatal cell types in spruce remains to be determined.

*pAtSPCH*-driven expression of *PaSM*, *PaFAMA*, *CrSM*, and *CrFAMA* in the *spch* background resulted in a loss of epidermal integrity. Cryo-scanning electron microscopy (Cryo-EM) images of *pAtSPCH:PaSM/spch* revealed defects resembling those observed in the *Arabidopsis bodyguard* (*bdg*) mutant^38^ (Supplementary Figure 3A). *BDG* has been proposed to contribute to the polymerization of carboxylic esters in either the cuticular layer of the cell wall or the cuticle proper^38,39^. This phenotypic similarity therefore raises the possibility that epidermal rupture in these transgenic lines involves defects in cuticle formation or in the organization of the cuticle–outer wall interface. However, defects in cuticle biosynthesis alone are unlikely to account for the loss of epidermal cells and exposure of underlying mesophyll tissue. We therefore propose that expression of *SM/FAMA* homologues in the *AtSPCH* domain disrupts the coordination between stomatal lineage differentiation with outer wall and cuticle assembly, thereby compromising epidermal integrity.

Beyond the comparative functional analysis, our anatomical observations provide additional context for stomatal development in Norway spruce. Like many gymnosperms and some tropical species, Norway spruce forms sunken stomata^25,40,41^. We observed that the GMC expands not only longitudinally and transversely but also along the adaxial–abaxial axis of the needle (Figure 1F), consistent with previous observations in pines^25^. Unlike in pine, however, Norway spruce stomata are not fully sunken immediately after GMC division, and the substomatal chamber remains only partially enclosed at this stage. These observations suggest that key developmental features—including asymmetric fate acquisition, GMC differentiation, and GC formation—are shared between Norway spruce and previously described conifers, while species-specific differences may exist in the timing of symmetric division and stomatal sinking. Such differences may reflect environmental influences or as-yet-unidentified regulatory mechanisms. As we could not find the un-sunken stomata formation in our spruce transgenic lines, its underlying mechanisms remain to be studied in future.

The limitation of this study should be acknowledged. Although the SRDX-fusion lines provide native-context evidence supporting roles for PaSM and PaFAMA in spruce stomatal development, definitive functional characterization in spruce will require loss-of-function approaches such as CRISPR-based mutagenesis or complementation under native regulatory sequences, since 1) we cannot remove repressed target by this technology so we can only see effects of removed upregulation, 2) our SRDX-fusion lines showed milder phenotype than *AtFAMA-EAR* which mimicked *fama* mutant phenotype in *Arabidopsis*^8^. The latter could be hypothetically caused by inefficient recruitment of chromatin remodeling components or histone acetylation modification enzyme by SRDX in spruce. An alternative explanation could be related to the promoter strength; since the promoter used in *Arabidopsis* line was inducible, it can potentially drive stronger expression than constitutive promoter used in spruce in this study. CRISPR lines remain technically challenging in gymnosperms but will be essential for establishing the precise developmental functions of these genes in their native context. Nevertheless, the convergence of cross-species complementation, expression patterns within spruce stomatal lineage, and native perturbation provides independent evidence demonstrating that these regulators are neither fully redundant nor as strictly specialized as the canonical angiosperm SPCH, MUTE, and FAMA system.

Together, our results show that broad developmental competence and functional specialization can coexist: the fern and gymnosperm bHLHs examined here act across multiple stomatal-lineage transitions while displaying distinct functional biases and, in spruce, they show partially divergent developmental functions strongly coinciding with their expression patterns. These findings support a model in which the pronounced stage specialization of angiosperm stomatal bHLHs emerged through progressive partitioning of broader, overlapping developmental capacities.

## Materials and Methods

### Phylogenetic analysis

The liverwort *Marchantia polymorpha*^42^, moss *Physcomitrium patens*^43^, lycophyte *Selaginella moellendorffii*^44^, ferns *Alsophila spinulosa*^45^ and *Ceratopteris richardii*^46^, gymnosperms *Ginkgo biloba*^47^*, Picea abies*^48^, *Pinus lambertiana*^49^, *Pseudotsuga menziesii*^50^ and *Sequoiadendron giganteum*^51^ and angiosperms *Arabidopsis thaliana*^52^, *Amborella trichopoda*^53^ *and Oryza sativa*^54^ were selected for phylogenetic analysis. Orthofinder^55^ (version 3.1.3) was run using amino acid sequences from the selected species obtained from Phytozome^56^, Fernbase (https://fernbase.org/), TreeGenes^57^, PlantGenIE^58^ and Ginkgo Database^59^ (Supplementary file 1). The orthogroup containing AtSPCH, AtMUTE and AtFAMA was selected as the bHLH1a orthogroup. The *Arabidopsis* bHLH3c genes *AT5G57150* and *AT4G2SS30* were included as outgroups.

Amino acid sequences were aligned using MAFFT (version 7.505)^60^ with the L-INS-i option. A phylogenetic tree was constructed with RAxML^61^ version 8.2.12. Based on the preliminary alignment and phylogenetic tree, the sequences looking potentially partial or otherwise misannotated gene models were manually re-annotated using FGENESH+^62^. The TBLASTN^63^ was used to identify unannotated genes and manual annotation was carried out with FGENESH+. The curated data (Supplementary file 2) was aligned with MAFFT with L-INS-i option. The resulting sequence alignment (Supplementary file 3) was used to construct a phylogenetic tree with RAxML using PROTGAMMAAUTO option to define the best-fit protein substation model, which for this data was JTT and - f a for tree estimation and bootstrapping with the 1000 bootstraps (Supplementary file 4). Tree was illustrated with FigTree (version 1.4.4) (https://github.com/rambaut/figtree/).

The amino acid sequences of AtFAMA, PaFAMA and CrFAMA were aligned using ClustalW^64^ with default settings in MEGA11^65^. The region corresponding to the previously described angiosperm FAMA-specific sequence^3^ was inspected in the alignment. To assess conservation of this region, the alignment columns corresponding to the FAMA-specific region in the reference angiosperm FAMA sequence were extracted from the full-length alignment. The extracted region was exported as an amino acid alignment and used for pairwise sequence comparison. Pairwise amino acid distances were calculated in MEGA11 using the p-distance model, with gaps and missing data treated by pairwise deletion. Percent amino acid identity was calculated as follows:

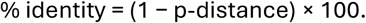

### Plant material and growth conditions

*Arabidopsis thaliana* Columbia-0 (Col-0) was used as the wild type, all transgenic lines and mutants were in the Col-0 genetic background. The *spch-3*, *mute-1*, *fama-1* were described previously^6–8^. Seeds of *pAtMUTE:AtMUTE-CFP* line were kindly provided from Dr. Dominique Bergmann.

*Arabidopsis* seeds were sown on half-strength Murashige and Skoog (MS) medium (Prod. No: M0221.0100, Duchefa Biochemie), adjusted to pH 5.75 and solidified with 0.8% plant agar (Prod. No: P1001.1000, Duchefa Biochemie). After stratification for at least two days at 4 °C, plates were incubated at 23 °C under long-day condition (16 h light/8 h dark).

Norway spruce seeds were obtained from the Natural Resources Institute Finland (Luke). One seed lot was used for temporal gene-expression analysis, whereas a separate seed lot was used for morphological analysis of stomatal complexes (Figure 1) and whole-needle imaging (Supplementary Figure 1A). A third seed lot was used for HCR RNA-FISH experiments.

Norway spruce seeds were sown on a mixture of peat and vermiculite (4:1) in plates and stratified at 4 °C for at least one week. The plates were then transferred to 20 °C under continuous light conditions. After germination, the seedlings were transferred to individual pots and supplied weekly with Neko fertilizer diluted 1:1,000 (KASVI-RAVINNE, NPK 7-2-2, Neko, (https://www.neko.fi/)). The day on which plates were transferred to the growth chamber was defined as day 0.

Surface-sterilized spores of *Ceratopteris richardii* were obtained from Carolina Biological Supply Company (https://www.carolina.com). Spores were suspended in sterile Milli-Q water and spread onto half-strength MS medium (Prod. No: M0221.0100, Duchefa Biochemie), adjusted to pH 5.75 and solidified with 0.8% plant agar (Prod. No: P1001.1000, Duchefa Biochemie) with spreader. Without stratification, the plates were incubated at 23 °C under long-day condition (16 h light/8 h dark). Approximately one-month-old gametophyte was moved to fresh half-strength MS medium and partially submerged in sterile Milli-Q water to promote fertilization. The day on which plates were transferred to the growth chamber was defined as day 0.

Proembryogenic masses (PEMs) of Norway spruce embryogenic cell line 11:12:04 initiated from seeds provided by the Swedish Forest Research Institute in 2011 were used for the spruce transformation. Prior to transformation, the cell line was propagated in vitro on semi-solid ½ LP medium supplemented with 10 µM 2,4-dichlorophenoxyacetic acid (2,4-D), 4 µM benzyl adenine (BA), 1% sucrose and 0.35% gelrite in darkness at 21-22°C and subcultured onto medium of the same composition every 2-3 weeks^66^.

### Plasmid construction and plant transformation

The coding sequence of *PaSM* (1383 bp) and *PaFAMA* (1428 bp) were amplified from cDNA prepared from frozen needles of *Picea abies* clone Z4006 (Köttsjö spruce). RNA was extracted as described below. The *CrSM* genomic sequence (2566 bp) was amplified from genomic DNA isolated from C.fern sporophyte leaves, as described in the genomic DNA extraction section.

All plasmids used for cross-species complementation analysis were generated utilizing Multisite Gateway Technology. For first-entry clone, *pAtSPCH* and *pAtMUTE* plasmids. *AtFAMA* promoter, containing approximately 3kb upstream of the start codon, was amplified from genomic DNA of *Arabidopsis* Col-0 using previously reported primers^7^ in pDONR221/Zeo. Second-entry clones contained the PaSM, PaFAMA coding sequences or *CrSM* genomic sequence without the stop codon in pDONR221/Zeo. The *CrFAMA* coding sequence (1077 bp) was synthesized by Biomatik (https://www.biomatik.com/) into pDONR221/Kan based on the publicly available *C. richardii* sequence Ceric.24G025300.1 (genome version 2.1; Phytozome 14 (https://phytozome-next.jgi.doe.gov/)). Third-entry clones contained triple Venus and the OCS terminator, p2R3a-3xVenusYFP-OcsT, was described in Siligato et al^67^. Multisite Gateway LR reactions were performed using pFRm43GW or pHm43GW as the destination vector.

Each expression vector was introduced into *Agrobacterium tumefaciens* by electroporation. The transformed *Agrobacterium* cultures were centrifuged, and the pellets were resuspended in a solution containing 5% sucrose, and 0.05% Silwet. The *Agrobacterium* suspension was applied to the shoot apical meristem of *Arabidopsis spch*, *mute*, or *fama* heterozygous mutants.

For spruce transformation, the hygromycin resistance gene HptII under the control of the maize ubiquitin promoter and the pea RbcS terminator was amplified from the p6oUZm-ActOs-Gusi vector (https://dna-cloning.com/binaries/) with the U078 and U077 primers flanked respectively with the F and G Greengate overhangs designed by Lampropoulos et al.^68^ (Supplemental Table 1, 2). The amplified PCR product was subsequently subcloned into the pGGF000. The PaSM (1383 bp) and PaFAMA (1428 bp) coding sequences were fused at their C termini to the SRDX repression domain in pGGC000 vector^31^. The *PaSM–SRDX* and *PaFAMA–SRDX* fragments were amplified from unpublished Multisite Gateway constructs containing *35S:PaSM–SRDX* and *35S:PaFAMA–SRDX*, respectively. Consequently, each fusion contained a 27-bp attL2-derived sequence between the coding sequence and the SRDX sequence. GreenGate assembly was used to place either *PaSM–SRDX* or *PaFAMA–SRDX* under the control of the CaMV 35S promoter and the RbcS terminator. Each expression cassette was assembled together with the HptII selection cassette into the pGGZ003 destination vector (Supplementary Table 2). The resulting constructs were introduced into Norway spruce by *Agrobacterium*-mediated transformation as described below (see “Agrobacterium-mediated Norway spruce transformation”).

### *Agrobacterium*-mediated Norway spruce transformation

^66^PEMs were prepared for *Agrobacterium* transformation 10-14 days after subculturing by resuspending 3 g of PEMs in 20ml liquid ½ LP medium containing 100 µM acetosyringone and then inoculated for 3 days on a rotary shaker at 110 rpm in the dark at 20-21°C. Unless otherwise specified, the pH of all media used for transformation and following plant production was adjusted to 5.8 before autoclavation.

A dense culture of *Agrobacterium tumefaciens* containing the vector with the gene of interest were scraped from LB agar plates containing the appropriate selection antibiotics and resuspended in infiltration buffer containing 10 mM MgCl_2_, 10 mM MES (pH 5.5), and 100 µM acetosyringone, then incubated for 1.5–3 h on a rotary shaker at 110 rpm in the dark at 20–21 °C.

The flask containing 20ml of resuspended PEMs was diluted 1:2 with ½ LP medium containing 100µM acetosyringone and the induced bacterial suspension to reach a final OD600 of 0.5 for 1-1.5h by co-cultivation on a rotary shaker at 110 rpm in darkness at 20–21 °C.

After co-cultivation, the PEMs suspension was filtered onto a paper disc (70 mm Whatman no 1 filterpaper, Cytiva) transferred to a 9-cm Petri dish with semi-solid ½ LP medium supplemented with 100 µM acetosyringone and incubated for 2 days in darkness at 21-22°C.

Selection of putatively transformed PEMs was performed on semi-solid ½ LP medium supplemented with 400 mg/L timentine and hygromycin in darkness at 21-22°C with subcultures every 2-3 weeks until growing clumps of PEMs appeared. The hygromycin concentration used for selection was 0.5 mg/L for the first 2 weeks followed by 2.5 mg/L. The number of PEMs-clumps that had reached 3-4 mm in diameter was recorded and the clumps isolated to be established as putatively transgenic cell lines on semi-solid ½ LP medium for further propagation and plant production. Putative transformed PEMs were verified by genotyping PCR and qRT-PCR as described below.

The verified transformed spruce embryogenic tissue was transferred from semi-solid ½ LP medium to semi-solid DKM maturation medium^66^ containing 3% sucrose, 0.5 g/L casein hydrolysate, 0.35% gelrite and cultivated for 7 days in the darkness at 21–22 °C. The cultures were then transferred to semi-solid DKM maturation medium supplemented with 29 µM ABA and cultured for 3 weeks, followed by subculturing onto fresh maturation medium supplemented with 29 µM ABA for an additional 3 weeks under the same conditions. Mature somatic embryos were manually picked and dried for 2 weeks under high humidity, in darkness at 21-22°C then germinated in squared 10 cm petri dishes on semi-solid AD1 germination medium^69^ containing 3% sucrose, 0.35% gelrite and 0.5 g/L casein hydrolysate. Petri plates with germinating somatic embryos were placed for one week in dark at 21-22°C followed by a constant red light (wavelength: 660 nm; TL-D 18W/15, Philips, Stockholm, Sweden) at 5 μmol/m^2^/s at 21-22°C for 1-2 weeks. The Petri plates with geminants were gradually moved to constant white, fluorescent light (Fluora L 18W/77, Osram, Johanneshov, Sweden) at 55-60 μmol/m^2^/s at 22°C until fully developed with root and shoot^69,70^. Developed germinants (after approximately 2 months under light) were acclimatized in a peat:perlite (80:20) composition under 24 h light conditions (Valoya LED lamps) at 110-120 μmol/m^2^/s at 21-22°C. The spruce plants were fertilized once per week using RIKA-s (SW, Horto).

### RNA extraction

For cloning of *Picea abies* genes, total RNA was extracted from approximately 100 mg of frozen tissue using a CTAB-based method modified from Chang et al^71^. Tissue was ground to a fine powder in liquid nitrogen and suspended in 750 µl of extraction buffer containing 2% (w/v) CTAB, 2% (w/v) PVP, 100 mM Tris–HCl (pH 8.0), 25 mM EDTA and 2 M NaCl, supplemented immediately before use with 2% (v/v) β-mercaptoethanol. Samples were incubated at 65°C for 1–3 min and extracted twice with an equal volume of chloroform:isoamyl alcohol (24:1, v/v). After centrifugation at 13,000 rpm for 10 min at room temperature, the aqueous phase was collected, and RNA was precipitated overnight at 4°C by adding one-quarter volume of 10 M LiCl. RNA was collected by centrifugation at 13,000 rpm for 15 min at 4°C and dissolved in nuclease-free water. RNA was further precipitated with 0.1 volume of 3 M sodium acetate and two volumes of absolute ethanol for at least 2 h at −20°C. The RNA pellet was washed with 70% ethanol, briefly air-dried and dissolved in nuclease-free water.

For qRT–PCR analysis of spruce needles, eight approximately one-month-old seedlings were pooled for each biological replicate. Needle samples were collected from four regions: the basal half of 0.2–0.3 cm needles, the basal half of 0.4–0.6 cm needles, the distal half of 0.4–0.6 cm needles, and the distal one-third of 0.8–0.9 cm needles. These samples were designated as stages 1, 2, 3, and 4, respectively. The predominant stomatal developmental stage in each sample group was determined by confocal microscopy. Stage 1 was characterized by prevalent ACDs, although some differentiating and dividing GMCs were also observed in this region (Figure 3A). Stage 2 consisted mainly of differentiating and dividing GMCs, with some ACDs also present (Figure 3B). Stage 3 was dominated by maturing stomata; GMC divisions were occasionally observed, whereas ACDs and GMC differentiation were absent (Figure 3C). Stage 4 consisted predominantly of maturing or mature GCs. For verifications of putative transformed spruce PEMs, an approximately 5 × 5 × 5 mm piece of proembryogenic mass was collected from each independently transformed line. RNA was extracted using modified versions of the CTAB protocol described above with reference to Pavy et al^72^. For needle samples, the extraction buffer was supplemented with 0.5 g l⁻¹ spermidine and 2% (v/v) β-mercaptoethanol. For proembryogenic mass samples, the buffer was supplemented with 0.5 g l⁻¹ spermidine, and β-mercaptoethanol was replaced with 2 M DTT added immediately before use at final concentration 40 mM. Frozen samples were ground in liquid nitrogen and mixed with 750 µl of extraction buffer preheated to 65°C. Samples were incubated at 65°C for 10 min, with intermittent vortexing, and extracted twice with 500 µl of chloroform:isoamyl alcohol (24:1, v/v). Following each extraction, samples were centrifuged at 21,000 × *g* for 5 min at room temperature, and the upper aqueous phase was transferred to a new tube. RNA was precipitated by adding an equal volume of 7.5 M LiCl containing 50 mM EDTA and incubating at −20°C for at least 1 h or overnight. RNA was collected by centrifugation at 21,000 × *g* at room temperature, washed with 80% ethanol, briefly air-dried and dissolved in sterile Milli-Q water.

### cDNA synthesis and qRT-PCR

RNA samples were treated with DNase I (Thermo Scientific, EN0521), and first-strand cDNA was synthesized using an oligo(dT)₁₈ primer and a First Strand cDNA Synthesis Kit (Thermo Scientific, K1612) according to the manufacturer’s instructions. qRT-PCR was performed using HOT FIREPol® EvaGreen® qPCR Mix Plus with passive reference dye with ROX (Solis BioDyne, 08-24-00001). For developmental-stage-specific expression analysis, qRT-PCR was performed using three biological replicates and three technical replicates. Expression levels were normalized to the internal control *Actin* and are presented relative to stage 1. Transgene expression in spruce transgenic lines was verified by qRT-PCR with one biological and technical replicate. Expression levels were normalized to the internal control *Actin* and are presented relative to WT. Relative gene expression levels were calculated using the 2^−ΔΔCt method.

### Genomic DNA extraction

C.fern sporophyte leaf (approximately 0.5×0.5 cm) was flash frozen with liquid nitrogen and was ground with frozen mortar and pestle. With the frozen material, we extracted DNA using DNeasy Plant Mini Kit (Qiagen, Cat no. / ID. 69104) following the provided protocol.

For verification of putatively transformed spruce PEMs, frozen PEM samples were allowed to thaw at room temperature for 5–10 min and suspended in 700 µl Edwards extraction buffer (200 mM Tris-HCl, pH 7.5, 250 mM NaCl, 25 mM EDTA, pH 8.0, and 0.5% SDS). The tissue was homogenised using a pestle. Samples were centrifuged at maximum speed for 10 min at room temperature, and 600 µl of the supernatant was transferred to a new tube containing an equal volume of isopropanol. DNA was precipitated overnight at −20 °C. The entire precipitated sample was transferred to a HiBind DNA Mini Column (DNACOL-01, Omega Bio-tek) and centrifuged at maximum speed for 30 s. Columns were washed twice with 700 µl DNA Wash Buffer containing ethanol (SPW Buffer, SPW-200, Omega Bio-tek), with centrifugation at maximum speed for 30 s after each wash, according to the manufacturer’s instructions. Columns were subsequently centrifuged empty at maximum speed for 2 min to remove residual ethanol. DNA was eluted by applying 50 µl elution buffer pre-warmed to 60 °C to the centre of the column, incubating for 2 min at room temperature, and centrifuging at maximum speed for 1 min. Extracted DNA was used as a template for genotyping PCR to test for Agrobacterium contamination and to confirm the presence of the *PaSM* or *PaFAMA* transgene (See Supplementary Table 1 for primers).

### Sample preparation and imaging

For differential interference contrast (DIC) microscopy imaging, plants were fixed in ethanol and acetic acid (7:1) and kept in fixative at least overnight. Samples were rinsed with Milli-Q water several times and mounted in Hoyer’s solution (7.5 g arabic gum, 5 ml glycerol, 103 g chloral hydrate and 30 ml Milli-Q water). After the tissues became transparent, cotyledons were observed using a Leica DM6B light microscope. 20×objective lens was used for imaging.

For confocal microscopy of *Arabidopsis* samples, seedlings were submerged in 0.1% (v/v) SCRI Renaissance Stain 2200 (SR2200) in 1× phosphate-buffered saline (PBS) using a centrifuge to stain cell wall. For propidium iodide (PI) staining, seedlings were submerged into 1 mg/mL PI solution for several seconds, followed by immediate wash with Milli-Q water. 20×objective lens was used for imaging.

For cross-species complementation analysis, unless otherwise indicated, all quantitative analysis and imaging were performed using plants homozygous for the respective mutant allele. T3 or T4 transgenic generations were used. Sample sizes are indicated in the corresponding plots.

Stomatal density was calculated as the number of stomata per unit epidermal area. Stomatal index was calculated as *SI* = *S*/(*S* + *E*) × 100, where *S*is the number of stomata [including/excluding abnormal stomata, as specified] and *E*is the number of other epidermal cells.

To describe wild-type spruce needle ontogeny, needles were excised from seedlings and fixed under vacuum in 4% paraformaldehyde (PFA) in 1×PBS containing 0.05% (v/v) Triton X-100. After one hour of vacuum treatment, the needles were washed several times with 1×PBS. For longitudinal imaging, the needles were transferred to 0.1% (v/v) SR2200 in ClearSee^73^. For cross-section imaging, needles were embedded in 4% agarose, and 200-um-thick cross-sections were prepared using a Leica Microm 650V vibratome. The sections were then transferred to 0.1% (v/v) SR2200 in ClearSee. 20×objective lens was used for imaging.

To analyze the morphology of transgenic spruce lines, seedlings were fixed under vacuum for approximately 15 minutes in 4% PFA in 1×PBS containing 0.05% (v/v) Triton X-100. Larger seedlings were cut into several pieces before fixation. After vacuum treatment, samples were stored in the fixative at 4 °C until preparation for observation. Needles were cut longitudinally halfway through with a razor blade under a stereomicroscope and immersed in 100 mM NaOH. Samples were then incubated at 65°C for 5–30 min, and rinsed several times with distilled water. Mesophyll cells and vascular tissues were removed as thoroughly as possible. The remaining needle epidermal tissues were immersed in 0.1% (v/v) SR2200 in ClearSee.

Stained samples were imaged with a Leica Stellaris 8 confocal microscope or a Leica DM6B fluorescence microscope. The number of epidermal cells separating adjacent stomata was quantified from stitched images over the longest continuous region in which each stomatal file could be reliably traced. For morphological analyzing spruce transgenic lines, z-stack images were acquired. the optical section in which the pore reached its maximum length was selected. For GC length and width measurements, the optical section in which the guard cells reached their maximum dimensions was selected. Pores and GCs were manually outlined in ImageJ, and pore length and GC length were measured as the maximum Feret diameter, whereas GC width was measured as the minimum Feret diameter. Sample sizes are provided in Supplementary Table 3. 20× or 10×objective lens was used for imaging.

### HCR RNA-FISH

HCR RNA-FISH was performed using the HCR™ RNA-FISH (v3.0) system (Molecular Instruments, https://www.molecularinstruments.com/) (Dirks and Pierce^74^; Choi et al.^75^). HCR™ probes (v3.0) targeting *PaSM* or *PaFAMA* were designed by Molecular Instruments using the standard number of binding sites (up to 20). For *PaSM* detection, the B4 amplifier conjugated to fluorophore 647 was used, whereas the B3 amplifier conjugated to fluorophore 546 was used for *PaFAMA* detection. Each *PaSM* and *PaFAMA* probe hybridization was performed separately. The protocol used in this study was adapted from Huang et al.^76^ and Oliva et al^77^, with some modifications.

Approximately 0.5-cm-long needles located close to the SAM were collected from 7-week-old Norway spruce seedlings grown from seeds provided by Luke. Needles were fixed in 4% PFA under vacuum for 10 min, followed by incubation on a tube rotator for 50 min. If needles remained floating after fixation, the samples were centrifuged for 1 min at maximum speed. Only fully submerged needles were used for subsequent steps.

After fixation, the needles were washed several times with 1× PBS and embedded in 5% agarose. The resulting agarose blocks were sectioned at 100 µm using a Leica Microm 650V vibratome. Sections were dehydrated sequentially in 70% ethanol, 90% ethanol, 100% ethanol twice and 100% methanol twice, with each step performed for 15 min. Samples were then incubated in methanol overnight at −20 °C.

The samples were rehydrated sequentially in 50% methanol/50% ethanol, 100% ethanol, 50% ethanol/50% PBS and PBST (1× PBS containing 0.1% Tween-20) twice, with each step performed for 10 min. Samples were then incubated with preheated HCR Probe Hybridization Buffer under vacuum for 10 min at room temperature, followed by pre-hybridization with gentle agitation for 1.5 h at 37 °C.

After pre-hybridization, the HCR Probe Hybridization Buffer was removed as completely as possible and replaced with probe solution containing 16 nM probe in HCR Probe Hybridization Buffer. Samples were incubated under vacuum for 10 min at room temperature and subsequently hybridized overnight (approximately 20 h) with gentle agitation at 37 °C. The probe solution was prepared by diluting a total of 8 pmol of probe in 500 µl HCR Probe Hybridization Buffer.

Following probe hybridization, the probe solution was removed as completely as possible. Samples were washed four times with preheated HCR Probe Wash Buffer at 37 °C for 15 min each, followed by two washes with 5× SSCT (5× saline-sodium citrate (SSC) containing 0.1% Tween-20) for 5 min each.

Samples were pre-incubated in HCR Amplification Buffer under vacuum for 10 min, followed by incubation with gentle agitation for 50 min. Hairpin solutions were prepared according to the manufacturer’s instructions. A total of 30 pmol of each hairpin was applied to the samples, which were first incubated under vacuum for 10 min and then incubated overnight (approximately 19 h) at room temperature.

Following amplification, samples were washed five times with 5× SSCT for 5 min, 5 min, 30 min, 30 min and 5 min, respectively. The wash buffer was removed as completely as possible, and samples were stained with SR2200 diluted 1:1,000 in ClearSee overnight or longer. Samples were imaged using a Leica Stellaris 8 confocal microscope. Z-stack images were acquired, and representative single optical sections containing clearly detectable RNA puncta were selected for presentation in Supplementary Figure 2.

### Immunolabeling and image analysis

Immunolabeling was performed largely as previously described^78^. Briefly, mature or maturing needles were excised from fixed seedlings and washed twice with 1× PBS. The needles were then embedded in 4% agarose dissolved in 1× PBS, and 200-µm-thick cross-sections were prepared using a Leica Microm 650V vibratome.

The sections were blocked in blocking buffer consisting of 2% (w/v) bovine serum albumin (BSA) in 1× PBS containing 0.1% (v/v) Tween-20 for at least 30 min at room temperature with gentle agitation. Primary antibodies, LM19 or LM20 (PlantProbes, University of Leeds), were diluted 1:200 in blocking buffer, and the samples were incubated in the diluted primary antibody solution overnight at 4°C. The next day, the samples were washed five times with blocking buffer.

Samples were then incubated overnight at 4°C with secondary antibody, anti-rat IgG Alexa Fluor 488, diluted 1:400 in blocking buffer. After incubation, the samples were washed five times with 1× PBS and cell walls were stained with 0.05% (v/v) SR2200 in 1× PBS. Samples were observed using a Leica Stellaris 8 confocal microscope.

Fluorescence signal intensities were quantified in Fiji by using the segmented line tool with spline fit to cover the signal of subsidiary cell wall at the interface with the guard cell. Mean gray value was measured and to compare signal intensities of the individual lines, values were normalized to the mean gray value of the corresponding antibiotic resistance control embedded in the same section.

### Cryo-EM

All cryo-SEM samples were prepared by means of the Quorum PP3010T Cryo-SEM preparation system. Firstly, the cut leaves were mounted on an aluminum stub by low temperature glue (Colloidal Graphite + O.T.C compound) and then plunged into slush nitrogen. After that, the deep-frozen samples were transferred into a pre-chamber at -140 °C, where a 5 nm thick platinum coating was deposited for reducing charging effect. Finally, the samples were loaded into the cryo-stage at 140 °C inside the JEOL JIB-4700F.

## Acknowledgments

We thank M. Lyytikäinen, M. Herpola, K. Kainulainen, I. Ahmad, M. Mäkelä, and H-L. Simola for technical assistance support, K. Nieminen, J. Alonso Serra and Vatén group members for comments and discussion. We thank Dr Dominique Bergmann at Stanford University and the Howard Hughes Medical Institute for providing multisite gateway compatible versions of *pAtMUTE* and *pAtSPCH* promoters and *spch-3*, *mute-1*, *fama-1*, and *pAtMUTE:AtMUTE-CFP* seeds, and Dr Ari Pekka Mähönen for providing triple Venus-OCST Multisite Gateway entry vector. We thank Dr Nathaniel Street from UPSC for providing frozen spruce needles and cDNA from P. abies clone Z4006 (Köttsjö spruce). We thank Natural Resources Institute Finland (Luke) for providing Norway spruce seeds. We would like to thank the gardeners L. Grönholm, D. Richterich. Confocal microscopy and histology experiment were performed at the Light Microscopy Unit and BI Histology Core Facility at the University of Helsinki, respectively. Transgenic spruce lines were generated in the Spruce Transformation Facility at Umeå Plant Science Centre. We acknowledge the provision of facilities for Cryo-SEM by Aalto University at the OtaNano-Nanomicroscopy Center (Aalto-NMC), especially L. Yao for technical help.

This research was funded by Research Council of Finland grant (Decision 340443), Research Council of Finland Centre of Excellence program 2022–2029 (Centre of Excellence in Tree Biology, Decisions 346140 and 364180), and University of Helsinki (ERC boost and HiLIFE Fellows program 2023-2025 grants to AV and DPPS fellowship to KE) and by the Wallenberg Initiatives in Forest Research (WIFORCE) funded by the Knut and Alice Wallenberg Foundation to ON and UE (KAW 2020.0240).

## Author Contributions

AV and MZI conceived and designed the research. MZI performed most of the experiments. KE contributed to the quantification of the cross-species complementation lines. AFMV performed phylogenetic analysis. VS performed the immunolabeling experiments and associated quantification. SG participated in experimental design. SJ, TD, UE, and ON optimized the spruce transformation protocol and generated the transgenic spruce lines. AV and MZI wrote the manuscript with input from all authors.

## Declaration of generative AI and AI-assisted technologies in the manuscript preparation process

During the preparation of this work, the authors used ChatGPT for language editing, including correcting grammar and improving clarity and readability. The authors reviewed and edited the output as needed and take full responsibility for the content of the published article.

---

**Supplementary Figure 1.**
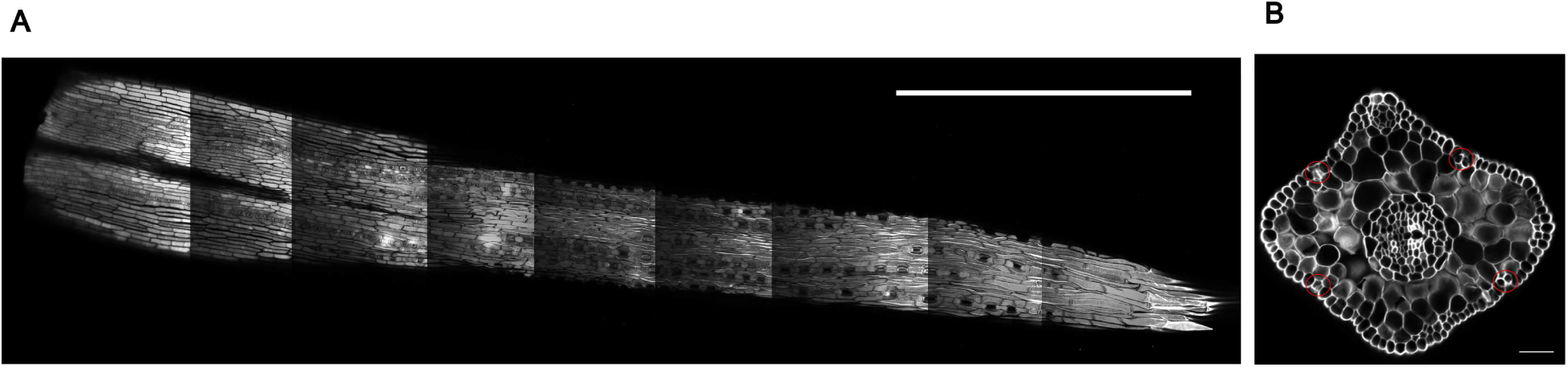
Spruce needles contain one to three stomatal files on each side. A, Longitudinal view of a young Norway spruce needle near the vegetative meristem. A 0.4 cm needle from a one-month-old seedling was imaged. **B,** Confocal image of cross-section of 1 cm needle from 4.25-month-old Norway spruce seedling. Red circles indicate the stomatal cells. Cell walls were stained with SR2200. Scale bar: 1 mm in **A**, 50 µm in **B**.

**Supplementary Figure 2.**
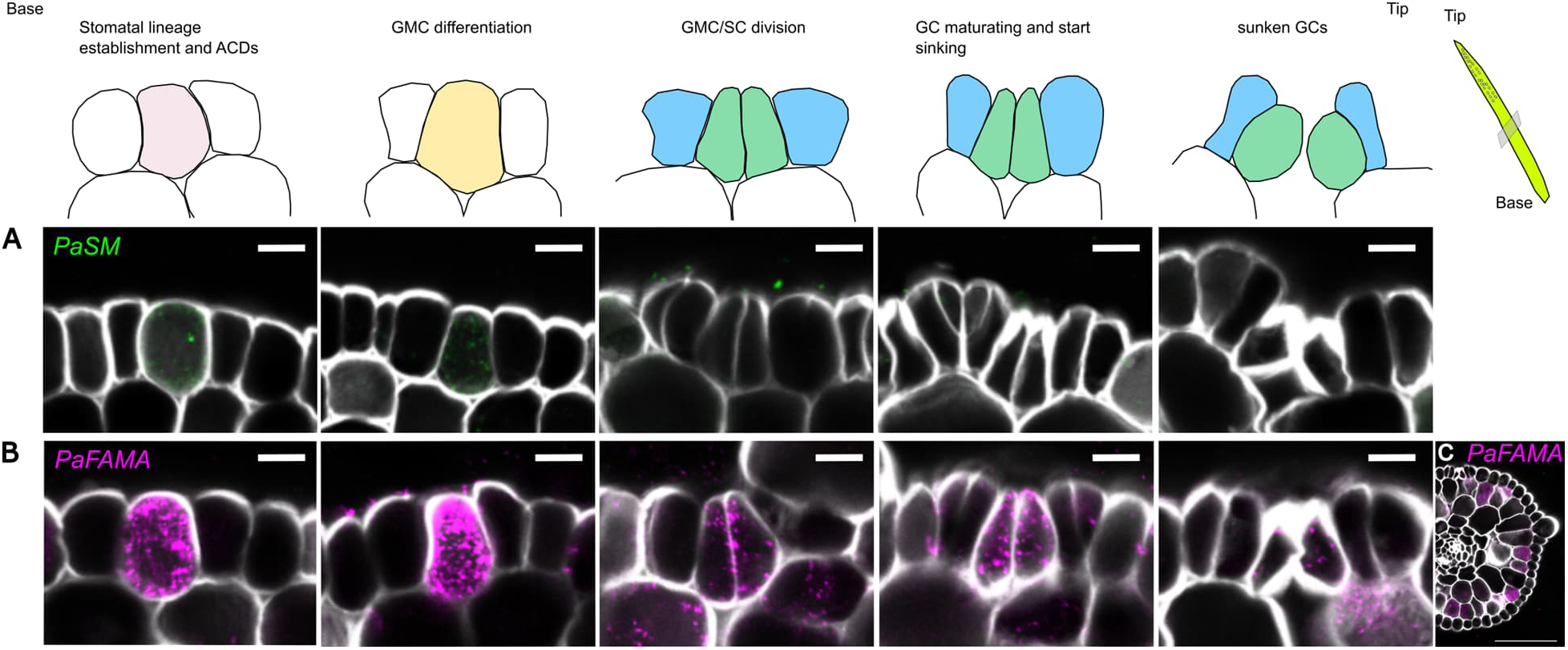
*PaSM* and *PaFAMA* show overlapping but distinct spatiotemporal expression patterns in the stomatal lineage. Schematic cross-sectional representation of stomatal development in a young Norway spruce needle near the SAM. Pink, yellow, green and blue indicate stomatal precursor cells, GMCs, GCs and lateral SCs, respectively. **A–C,** HCR RNA-FISH was performed on young Norway spruce needles using probes targeting *PaSM* and *PaFAMA*. The experiment was performed once using nine needles collected from three individual seedlings. **A,** *PaSM* transcripts were detected in early stomatal-lineage cells and during GMC differentiation but were not observed at later stages of stomatal development. **B,** *PaFAMA* transcripts were detected throughout multiple stages of stomatal development, including early stomatal-lineage cells, differentiating and dividing GMCs, and variably in mature GCs. **C,** PaFAMA trasncripts were detected in mesophyll cells. Cell walls were stained with SR2200. Scale bar, 10 µm (**A**, **B**), 100 µm (**C**).

**Supplementary Figure 3.**
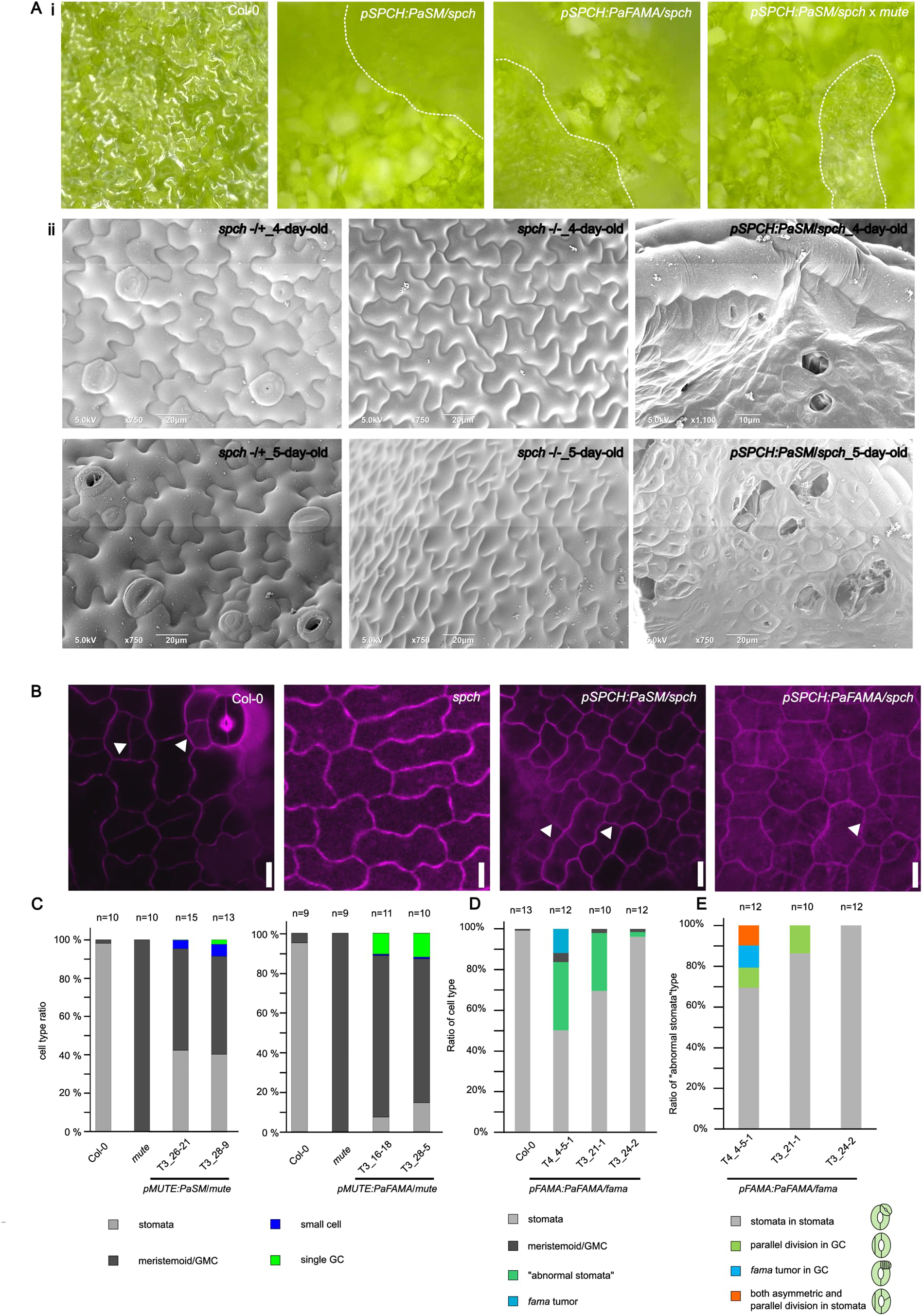
*PaSM* and *PaFAMA* expression in *Arabidopsis* stomatal mutants causes diverse stomatal and epidermal phenotypes. **A**-**i**, The abaxial surface of 10-day-old cotyledons of Col-0, *pAtSPCH:PaSM/spch-3*, *pAtSPCH:PaFAMA/spch-3*, and *pAtSPCH:PaSM/spch-3 × mute* were imaged by microscopy camera, Dino-Lite microscope. The *spch* mutant genotype was not determined in *pAtSPCH:PaFAMA/spch-3,* whereas the other lines were homozygous for the *spch* mutation. The *mute* mutant background was homozygous in *pAtSPCH:PaSM/spch-3 × mute.* **A-ii,** Cryo-EM images of 4-day-old or 5-day old abaxial cotyledons of *spch-/+*, *spch-/-*, and *pAtSPCH:PaSM/spch-3*. **B**, Confocal images of 2-day-old abaxial cotyledons of Col-0, *spch-3* mutant, *pAtSPCH:PaSM/spch-3,* and *pAtSPCH:PaFAMA/spch-3.* In *pAtSPCH:CrFAMA* lines, most epidermal cells showed intracellular signal with propidium iodide, possibly due to temporary changes in membrane permeability, differences in cell wall, membrane structure, cuticule, or cell death. **C**, Ratio of cell types at 10-day-old abaxial cotyledon of *pAtMUTE:PaSM/mute-1* and *pAtMUTE:PaFAMA/mute-1.* **D**, Ratio of cell types at 10-day-old abaxial cotyledon of *pAtFAMA:PaFAMA/fama-1.* **E**, Ratio of cell types of “abnormal stomata” at 10-day-old abaxial cotyledon of *pAtFAMA:PaFAMA/fama-1.* White dashed lines indicate the boundary between epidermal and mesophyll cells. Magenta, propidium iodide. Scale bar, 10 µm in **B**.

**Supplementary Figure 4.**
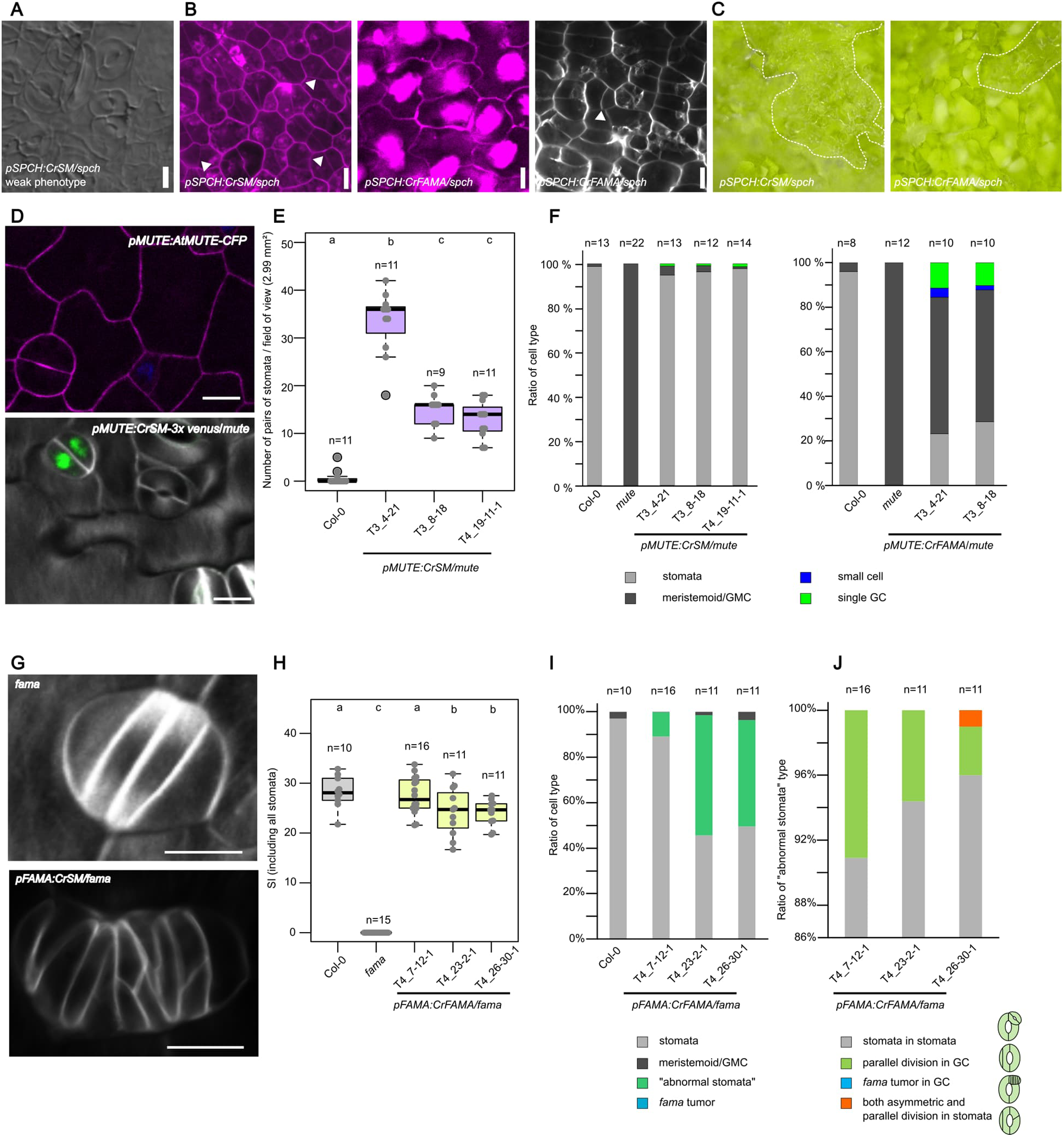
Additional characterization of stomatal phenotypes in *Arabidopsis* lines complemented by *CrSM* and *CrFAMA*. **A**, DIC images of 10-day-old abaxial cotyledon of *pAtSPCH:CrSM/spch-3* weak phenotype line. Stomatal clusters were observed. **B**, Confocal images of the abaxial cotyledon epidermis of 2-day-old *pAtSPCH:CrSM/spch-3* and *pAtSPCH:CrFAMA/spch-3.* **C**, Microscope camera images acquired using a Dino-Lite microscope showing the abaxial surface of 10-day-old cotyledons of *pAtSPCH:CrSM/spch-3* and *pAtSPCH:CrFAMA/spch-3*. The *spch* mutant background was homozygous. **D**, Confocal images of abaxial cotyledons from 2-day-old *pAtMUTE:AtMUTE-CFP* seedlings and 10-day-old *pAtMUTE:CrSM-3x venus/mute-1* seedlings. **E,** Number of pairs of stomata at 10-day-old abaxial cotyledon of *pAtMUTE:CrSM /mute-1.* **F**, Ratio of cell types at 10-day-old abaxial cotyledon of *pAtMUTE:CrSM/mute-1* and *pAtMUTE:CrFAMA/mute-1.* **G,** Confocal images of *fama* tumors in abaxial cotyledons from 10-day-old *fama-1* and *pAtFAMA:CrSM/fama-1* seedlings. **H,** SI on the abaxial side of cotyledons in 10-day-old *pAtFAMA:CrFAMA/fama-1* seedlings. SI was calculated using both normal stomata and “abnormal stomata”. **I**, Ratio of cell types at 10-day-old abaxial cotyledon of *pAtFAMA:CrFAMA/fama-1.* **J**, Ratio of cell types of stomata with extra divisions at 10-day-old abaxial cotyledon of *pAtFAMA:CrFAMA/fama-1.* For **E** and **H,** statistical significance was assessed by the Steel-Dwass-Critchlow-Fligner test; different letters indicate statistically significant differences between groups (*p* < 0.05). Box plots show the median and interquartile range, with whiskers extending to values within 1.5× the interquartile range. Individual data points are shown as beeswarm plots. Cell walls were stained with SR2200 (**B**,**D**,**G**), except where shown in magenta, which indicates propidium iodide staining. Scale bar, 10 µm.

**Supplementary Figure 5.**
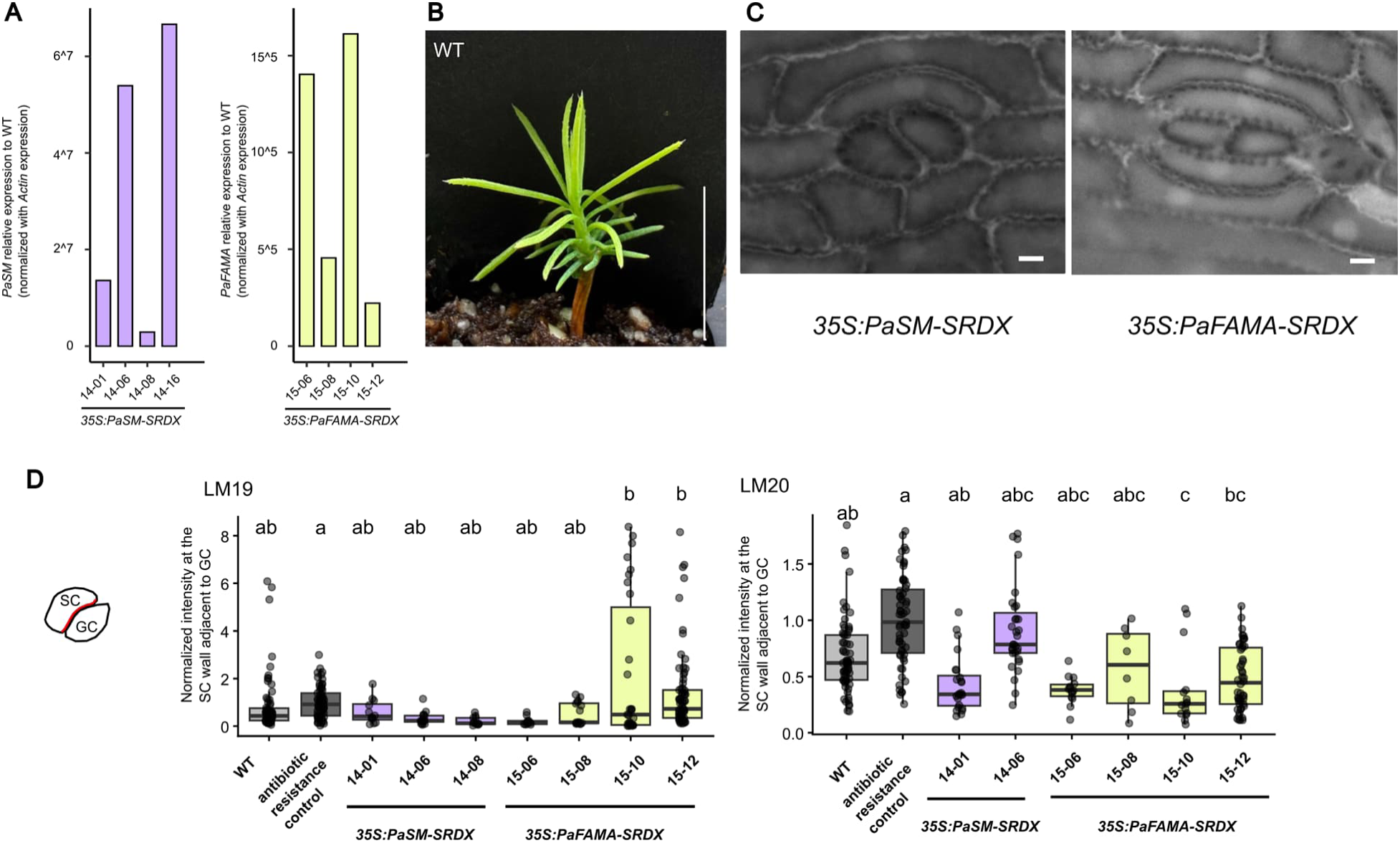
*PaSM* and *PaFAMA* alter aspects of spruce stomatal development. **A**, qRT-PCR analysis of *PaSM* and *PaFAMA* expression levels in proembryogenic mass samples from *35S:PaSM-SRDX* and *35S:PaFAMA-SRDX* lines, respectively. **B**, One-month-old wild type seedling. **C**, Fluorescence microscopy image of spruce transgenic lines showing potentially aborted GMCs. **D**, Quantification of LM19 and LM20 signal intensity in the SC wall at the interface with GC. Statistical significance was assessed using a linear mixed-effects model followed by pairwise comparisons (Tukey method, no additional adjustments). Different letters indicate statistically significant differences between groups (*P* < 0.05). Scale bar, 1 cm (**B**), 10 µm (**C**).

**Supplementary file 1 Information of the data origin and curation for species used in the phylogenetic analysis**

**Supplementary file 2 Amino acid sequences used in phylogenetic analysis in fasta format Supplementary file 3 Alignment of the sequences in fasta format**

**Supplementary file 4 Phylogenetic tree in NEXUS format with bootstrapping values**

**Supplementary Table 1 List of primers**

**Supplementary Table 2 List of vectors for spruce transformation**

**Supplementary Table 3 Number of samples observed for spruce transgenic analysis**

## Reference

1. Hetherington, A. M. & Woodward, F. I. The role of stomata in sensing and driving environmental change. Nature 424, 901–908 (2003).

2. Vatén, A. & Bergmann, D. C. Mechanisms of stomatal development: an evolutionary view. EvoDevo 3, 11 (2012).

3. MacAlister, C. A. & Bergmann, D. C. Sequence and function of basic helix-loop-helix proteins required for stomatal development in Arabidopsis are deeply conserved in land plants: Conservation of stomatal bHLHs. Evolution & Development 13, 182–192 (2011).

4. Nunes, T. D. G., Zhang, D. & Raissig, M. T. Form, development and function of grass stomata. The Plant Journal 101, 780–799 (2020).

5. Raissig, M. T. et al. Mobile MUTE specifies subsidiary cells to build physiologically improved grass stomata. Science 355, 1215–1218 (2017).

6. MacAlister, C. A., Ohashi-Ito, K. & Bergmann, D. C. Transcription factor control of asymmetric cell divisions that establish the stomatal lineage. Nature 445, 537–540 (2007).

7. Pillitteri, L. J., Sloan, D. B., Bogenschutz, N. L. & Torii, K. U. Termination of asymmetric cell division and differentiation of stomata. Nature 445, 501–505 (2007).

8. Ohashi-Ito, K. & Bergmann, D. &. *Arabidopsis* FAMA Controls the Final Proliferation/Differentiation Switch during Stomatal Development. The Plant Cell 18, 2493–2505 (2006).

9. Kanaoka, M. M., et al. *SCREAM/ICE1* and *SCREAM2* Specify Three Cell-State Transitional Steps Leading to *Arabidopsis* Stomatal Differentiation. The Plant Cell 20, 1775–1785 (2008).

10. Ortega, A., De Marcos, A., Illescas-Miranda, J., Mena, M. & Fenoll, C. The Tomato Genome Encodes SPCH, MUTE, and FAMA Candidates That Can Replace the Endogenous Functions of Their Arabidopsis Orthologs. Front. Plant Sci. 10, 1300 (2019).

11. Liu, T., Ohashi-Ito, K. & Bergmann, D. C. Orthologs of *Arabidopsis thaliana* stomatal bHLH genes and regulation of stomatal development in grasses. Development 136, 2265–2276 (2009).

12. McKown, K. H. et al. Expanded roles and divergent regulation of FAMA in Brachypodium and Arabidopsis stomatal development. The Plant Cell 35, 756–775 (2023).

13. Raissig, M. T., Abrash, E., Bettadapur, A., Vogel, J. P. & Bergmann, D. C. Grasses use an alternatively wired bHLH transcription factor network to establish stomatal identity. Proc. Natl. Acad. Sci. U.S.A. 113, 8326–8331 (2016).

14. Spiegelhalder, R. P. et al. Dual role of BdMUTE during stomatal development in the model grass Brachypodium distachyon. Development 151, dev203011 (2024).

15. Cheng, X. et al. MUTE drives asymmetric divisions to form stomatal subsidiary cells in Crassulaceae succulents. Sci. Adv. 12, eaeb8145 (2026).

16. Song, Z. et al. EgSPEECHLESS Responses to Salt Stress by Regulating Stomatal Development in Oil Palm. IJMS 23, 4659 (2022).

17. Lopez-Anido, C. B. et al. Single-cell resolution of lineage trajectories in the Arabidopsis stomatal lineage and developing leaf. Developmental Cell 56, 1043–1055.e4 (2021).

18. González-Suárez, P. et al. Targets of SPEECHLESS and FAMA control guard cell division and expansion in the late stomatal lineage. Development 153, dev205374 (2026).

19. Ran, J.-H., Shen, T.-T., Liu, W.-J. & Wang, X.-Q. Evolution of the bHLH Genes Involved in Stomatal Development: Implications for the Expansion of Developmental Complexity of Stomata in Land Plants. PLoS ONE 8, e78997 (2013).

20. Harris, B. J., Harrison, C. J., Hetherington, A. M. & Williams, T. A. Phylogenomic Evidence for the Monophyly of Bryophytes and the Reductive Evolution of Stomata. Current Biology 30, 2001–2012.e2 (2020).

21. Davies, K. A. & Bergmann, D. C. Functional specialization of stomatal bHLHs through modification of DNA-binding and phosphoregulation potential. Proc. Natl. Acad. Sci. U.S.A. 111, 15585–15590 (2014).

22. Chater, C. C. et al. Origin and function of stomata in the moss Physcomitrella patens. Nature Plants 2, 16179 (2016).

23. Moriya, K. C. et al. Stomatal regulators are co-opted for seta development in the astomatous liverwort Marchantia polymorpha. Nat. Plants 9, 302–314 (2023).

24. Chang, G. et al. Liverwort bHLH transcription factors and the origin of stomata in plants. Current Biology 33, 2806–2813.e6 (2023).

25. Johnson, R. W. & Riding, R. T. STRUCTURE AND ONTOGENY OF THE STOMATAL COMPLEX IN PINUS STROBUS L. AND PINUS BANKSIANA LAMB. American J of Botany 68, 260–268 (1981).

26. Toledo-Ortiz, G., Huq, E. & Quail, P. H. The Arabidopsis Basic/Helix-Loop-Helix Transcription Factor Family[W]. The Plant Cell 15, 1749–1770 (2003).

27. Heim, M. A. The Basic Helix-Loop-Helix Transcription Factor Family in Plants: A Genome-Wide Study of Protein Structure and Functional Diversity. Molecular Biology and Evolution 20, 735–747 (2003).

28. Lampard, G. R., MacAlister, &. A. C Bergmann, D. C. *Arabidopsis* Stomatal Initiation Is Controlled by MAPK-Mediated Regulation of the bHLH SPEECHLESS. Science 322, 1113–1116 (2008).

29. Matos, J. L. et al. Irreversible fate commitment in the Arabidopsis stomatal lineage requires a FAMA and RETINOBLASTOMA-RELATED module. eLife 3, e03271 (2014).

30. Mahoney, A. K. et al. Functional analysis of the Arabidopsis thaliana MUTE promoter reveals a regulatory region sufficient for stomatal-lineage expression. Planta 243, 987–998 (2016).

31. Hiratsu, K., Matsui, K., Koyama, T. & Ohme-Takagi, M. Dominant repression of target genes by chimeric repressors that include the EAR motif, a repression domain, in Arabidopsis. The Plant Journal 34, 733–739 (2003).

32. Davies, K. A. & Bergmann, D. C. Functional specialization of stomatal bHLHs through modification of DNA-binding and phosphoregulation potential. Proc. Natl. Acad. Sci. U.S.A. 111, 15585–15590 (2014).

33. Han, S.-K. et al. MUTE Directly Orchestrates Cell-State Switch and the Single Symmetric Division to Create Stomata. Developmental Cell 45, 303–315.e5 (2018).

34. Wallner, E.-S. et al. Spatially resolved proteomics of the Arabidopsis stomatal lineage identifies polarity complexes for cell divisions and stomatal pores. Developmental Cell 59, 1096–1109.e5 (2024).

35. Shirakawa, M. et al. FAMA Is an Essential Component for the Differentiation of Two Distinct Cell Types, Myrosin Cells and Guard Cells, in *Arabidopsis*. The Plant Cell 26, 4039–4052 (2014).

36. Zhang, C. et al. The Arabidopsis F-box protein FBS associated with the helix-loop-helix transcription factor FAMA involved in stomatal immunity. Plant Mol Biol 115, 48 (2025).

37. Franceschi, V. R., Krokene, P., Christiansen, E. & Krekling, T. Anatomical and chemical defenses of conifer bark against bark beetles and other pests. New Phytologist 167, 353–376 (2005).

38. Kurdyukov, S. et al. The Epidermis-Specific Extracellular BODYGUARD Controls Cuticle Development and Morphogenesis in *Arabidopsis*. The Plant Cell 18, 321–339 (2006).

39. Jakobson, L. et al. BODYGUARD is required for the biosynthesis of cutin in Arabidopsis. New Phytologist 211, 614–626 (2016).

40. Šantrůček, J. The why and how of sunken stomata: does the behaviour of encrypted stomata and the leaf cuticle matter? Annals of Botany 130, 285–300 (2022).

41. Rudall, P. J., Rowland, A. & Bateman, R. M. Ultrastructure of Stomatal Development in *Ginkgo biloba*. International Journal of Plant Sciences 173, 849–860 (2012).

42. Bowman, J. L. et al. Insights into Land Plant Evolution Garnered from the Marchantia polymorpha Genome. Cell 171, 287–304.e15 (2017).

43. Bi, G. et al. Near telomere-to-telomere genome of the model plant Physcomitrium patens. Nat. Plants 10, 327–343 (2024).

44. Banks, J. A. et al. The Selaginella Genome Identifies Genetic Changes Associated with the Evolution of Vascular Plants. Science 332, 960–963 (2011).

45. Huang, X. et al. The flying spider-monkey tree fern genome provides insights into fern evolution and arborescence. Nat. Plants 8, 500–512 (2022).

46. Marchant, D. B. et al. The C-Fern (Ceratopteris richardii) genome: insights into plant genome evolution with the first partial homosporous fern genome assembly. Sci Rep 9, 18181 (2019).

47. Liu, H. et al. The nearly complete genome of Ginkgo biloba illuminates gymnosperm evolution. Nat. Plants 7, 748–756 (2021).

48. Nystedt, B. et al. The Norway spruce genome sequence and conifer genome evolution. Nature 497, 579–584 (2013).

49. Stevens, K. A. et al. Sequence of the Sugar Pine Megagenome. Genetics 204, 1613–1626 (2016).

50. Neale, D. B. et al. The Douglas-Fir Genome Sequence Reveals Specialization of the Photosynthetic Apparatus in Pinaceae. G3 Genes|Genomes|Genetics 7, 3157–3167 (2017).

51. Scott, A. D. et al. A Reference Genome Sequence for Giant Sequoia. G3 Genes|Genomes|Genetics 10, 3907–3919 (2020).

52. Cheng, C. et al. Araport11: a complete reannotation of the *Arabidopsis thaliana* reference genome. The Plant Journal 89, 789–804 (2017).

53. Amborella Genome Project et al. The *Amborella* Genome and the Evolution of Flowering Plants. Science 342, 1241089 (2013).

54. Ouyang, S. et al. The TIGR Rice Genome Annotation Resource: improvements and new features. Nucleic Acids Research 35, D883–D887 (2007).

55. Emms, D. M., Liu, Y., Belcher, L., Holmes, J. & Kelly, S. OrthoFinder: scalable phylogenetic orthology inference for comparative genomics. Preprint at 10.1101/2025.07.15.664860 (2025).

56. Goodstein, D. M. et al. Phytozome: a comparative platform for green plant genomics. Nucleic Acids Research 40, D1178–D1186 (2012).

57. Wegrzyn, J. L., Lee, J. M., Tearse, B. R. & Neale, D. B. TreeGenes: A Forest Tree Genome Database. International Journal of Plant Genomics 2008, 1–7 (2008).

58. Sundell, D. et al. The Plant Genome Integrative Explorer Resource: PlantGen IE .org. New Phytologist 208, 1149–1156 (2015).

59. Gu, K.-J., Lin, C.-F., Wu, J.-J. & Zhao, Y.-P. GinkgoDB: an ecological genome database for the living fossil, Ginkgo biloba. Database 2022, baac046 (2022).

60. Katoh, K. MAFFT: a novel method for rapid multiple sequence alignment based on fast Fourier transform. Nucleic Acids Research 30, 3059–3066 (2002).

61. Stamatakis, A. RAxML version 8: a tool for phylogenetic analysis and post-analysis of large phylogenies. Bioinformatics 30, 1312–1313 (2014).

62. Solovyev, V. V. Statistical approaches in Eukaryotic gene prediction. in *Handbook of Statistical Genetics* 1616p. (Wiley-Interscience, 2007).

63. Camacho, C. et al. BLAST+: architecture and applications. BMC Bioinformatics 10, 421 (2009).

64. Thompson, J. D., Higgins, D. G. & Gibson, T. J. CLUSTAL W: improving the sensitivity of progressive multiple sequence alignment through sequence weighting, position-specific gap penalties and weight matrix choice. Nucl Acids Res 22, 4673–4680 (1994).

65. Tamura, K., Stecher, G. & Kumar, S. MEGA11: Molecular Evolutionary Genetics Analysis Version 11. Molecular Biology and Evolution 38, 3022–3027 (2021).

66. Arnold, S. V. & Clapham, D. Spruce Embryogenesis. in Plant Embryogenesis (eds Suárez, M. F. C Bozhkov, P. V.) vol. 427 31–47 (Humana Press, Totowa, NJ, 2008).

67. Siligato, R. et al. MultiSite Gateway-Compatible Cell Type-Specific Gene-Inducible System for Plants. Plant Physiol. 170, 627–641 (2016).

68. Lampropoulos, A. et al. GreenGate - A Novel, Versatile, and Efficient Cloning System for Plant Transgenesis. PLoS ONE 8, e83043 (2013).

69. Dobrowolska, I., Businge, E., Abreu, I. N., Moritz, T. & Egertsdotter, U. Metabolome and transcriptome profiling reveal new insights into somatic embryo germination in Norway spruce (Picea abies). Tree Physiology 37, 1752–1766 (2017).

70. Kvaalen, H. & Appelgren, M. Light quality influences germination, root growth and hypocotyl elongation in somatic embryos but not in seedlings of norway spruce. In Vitro Cell.Dev.Biol.-Plant 35, 437–441 (1999).

71. Chang, S., Puryear, J. & Cairney, J. A simple and efficient method for isolating RNA from pine trees. Plant Mol Biol Rep 11, 113–116 (1993).

72. Pavy, N. et al. Identification of conserved core xylem gene sets: conifer cDNA microarray development, transcript profiling and computational analysis. New Phytologist 180, 766–786 (2008).

73. Kurihara, D., Mizuta, Y., Sato, Y. & Higashiyama, T. ClearSee: a rapid optical clearing reagent for whole-plant fluorescence imaging. Development dev.127613 (2015) doi:10.1242/dev.127613.

74. Dirks, R. M. & Pierce, N. A. Triggered amplification by hybridization chain reaction. Proc. Natl. Acad. Sci. U.S.A. 101, 15275–15278 (2004).

75. Choi, H. M. T. et al. Third-generation *in situ* hybridization chain reaction: multiplexed, quantitative, sensitive, versatile, robust. Development 145, dev165753 (2018).

76. Huang, T., Guillotin, B., Rahni, R., Birnbaum, K. D. & Wagner, D. A rapid and sensitive, multiplex, whole mount RNA fluorescence in situ hybridization and immunohistochemistry protocol. Plant Methods 19, 131 (2023).

77. Oliva, M. et al. An environmentally responsive transcriptional state modulates cell identities during root development. Preprint at 10.1101/2022.03.04.483008 (2022).

78. Ye, L. et al. Cambium LBDs promote radial growth by regulating PLL-mediated pectin metabolism. Nat. Plants 11, 2565–2580 (2025).

79. Yang, X., Gavya S, L., Zhou, Z., Urano, D. & Lau, O. S. Abscisic acid regulates stomatal production by imprinting a SnRK2 kinase–mediated phosphocode on the master regulator SPEECHLESS. Sci. Adv. 8, eadd2063 (2022).

